# Tumor cell death by ferroptosis contributes to an immunosuppressive tumor microenvironment in syngeneic murine models of cancer

**DOI:** 10.64898/2026.01.09.698662

**Authors:** Nneka E. Mbah, Damien Sutton, Hanna S. Hong, Rashi Singhal, Rosa E. Menjivar, Heather Giza, Peter Sajjakulnukit, Matthew Perricone, Zeribe C. Nwosu, Jonathan Alektiar, Jason Lin, Daniel Long, Anthony C. Andren, Li Zhang, Howard C. Crawford, Timothy L. Frankel, Marina Pasca di Magliano, Luigi Franchi, Yatrik M. Shah, Costas A. Lyssiotis

**Affiliations:** Department of Molecular & Integrative Physiology, University of Michigan, Ann Arbor, MI, USA; Department of Internal Medicine, Division of Gastroenterology and Hepatology, University of Michigan, Ann Arbor, MI, USA; Rogel Cancer Center, University of Michigan, Ann Arbor, MI, USA; Graduate Program in Immunology, University of Michigan, Ann Arbor, MI, USA; Graduate Program in Cancer Biology, University of Michigan, Ann Arbor, MI, USA; Cellular and Molecular Biology program, University of Michigan, Ann Arbor, MI, USA; Department of Surgery, University of Michigan Medical School, Ann Arbor, MI, USA; Department of Pediatrics, University of Michigan Medical School, Ann Arbor, USA; Rogel and Blondy Center for Pancreatic Cancer, Rogel Cancer Center, University of Michigan, Ann Arbor, MI, USA

## Abstract

Pancreatic ductal adenocarcinoma (PDAC) is characterized by profound metabolic rewiring and a strongly immunosuppressive tumor microenvironment, both of which contribute to poor therapeutic responses. Immunogenic cell death (ICD) represents a potential strategy to overcome immune suppression by coupling tumor cell death to anti-tumor immune activation. Here, we investigated whether targeting amino acid metabolism in PDAC can induce ICD and promote tumor immunity. Through a focused metabolic screen in a panel of syngeneic mouse cancer cell lines, we identified cysteine restriction as a robust inducer of multiple damage-associated molecular patterns (DAMPs) in vitro, hallmark features of ICD. In addition to driving DAMPs, cystine-deprived tumor cells also promoted dendritic cell phagocytosis, maturation, and proinflammatory cytokine production in vitro. Because cysteine deprivation is a known trigger of ferroptosis, we further demonstrated that pharmacologic inhibition of glutathione peroxidase 4 (GPX4) similarly elicited ICD-associated features, which were reversible by the ferroptosis inhibitor Ferrostatin-1.

To define additional immune-modulatory signals associated with ferroptosis, we performed metabolomic and lipidomic profiling of cells undergoing, but not yet committed to, ferroptotic death. These analyses revealed selective release of immunosuppressive metabolites and oxidized phospholipids. Consistent with this, conditioned media from ferroptotic cells impaired CD8⁺ T cell proliferation and cytotoxicity in vitro. Thus, together our results indicated that the induction of ferroptotic immunogenic cell death led to the release of both pro- and anti-inflammatory signals. Subsequent analysis in vivo revealed that ferroptotic tumor cells predominantly contributed to a tumor-protective environment. In particular, tumors inoculated with ferroptotic cells were enriched with immunosuppressive myeloid cells and exhibited reduced populations of tumor-infiltrating CD8+ T cells. Further investigation using immune compromised mice suggested that ferroptotic cells may suppress both adaptive and innate immune responses. Collectively, these results underscore the complex and highly context-dependent effects of ferroptosis on tumor immunity, highlighting the critical importance of *in vivo* models to determine true immunogenic potential within the tumor microenvironment.

## Background

Pancreatic cancer is the third leading cause of cancer mortality in the United States^1,2^. Over 90% of these pancreatic malignancies are pancreatic ductal adenocarcinoma (PDAC), for which the overall 5-year survival rate is 13%^3^. Incidence of PDAC has been steadily increasing over the past two decades and it is projected to become the second-leading cause of cancer-related death by 2030^4^.

Challenges in treating PDAC result in part from features of the tumor microenvironment (TME)^5,6^. Within the TME lies an array of stromal cells that deposit a dense extracellular matrix, which leads to high intratumoral pressure and vascular collapse^7^. Consequences of hypovascularity are poor drug penetration, reduced nutrient delivery, and hypoxia^8–10^. Under these nutrient deregulated and oxygen depleted conditions, Kras mutations help to rewire metabolism in PDAC cells^11–13^. This metabolic reshaping involves increased nutrient recycling and scavenging, as well as metabolic crosstalk among cells in the TME^14–20^. Such metabolic adaptations not only support tumor survival but also undermine immune function by depleting nutrients essential for cytotoxic T cells and generating anti-inflammatory metabolites^21,22^. Immune function in the TME is further impaired by the presence of regulatory T cells, myeloid-derived suppressor cells, and tumor-associated macrophages, which act collectively to secrete factors that impede antitumor immunity^23^. Together, these architectural, metabolic, and cellular elements suppress anti-tumor immune responses and promote immune evasion^24^.

Recent advances in immune-based therapies are transforming treatment for many cancers by providing durable anti-tumor responses^25–28^. These innovations highlight the importance of engaging the immune system as an effective strategy against tumors. Immunogenic cell death (ICD) is a type of cell death that alerts the immune system by releasing immune stimulatory signals. Thus, inducing immunogenic cell death in cancer cells may offer a therapeutic approach to overcome the immunosuppressive TME of PDAC. Indeed, the immunogenic properties of various cancer therapies have been shown to synergize with immune checkpoint inhibitors and other immunomodulatory agents, both in clinical and preclinical studies, by increasing tumor antigen visibility and mitigating the immunosuppressive milieu of the TME^29–39^.

For cell death to be immunogenic, it must provide adjuvants—signals that activate the immune system—and be antigenic, resulting in the production of recognizable tumor antigens^40–42^. Therapies that induce ICD facilitate the release of adjuvants called damage-associated molecular patterns (DAMPs) from dying tumor cells^43^. Among the most consistently reported adjuvants produced during cell death are the DAMPs ecto-calreticulin, adenosine triphosphate (ATP), and High Mobility Group Box 1 (HMGB1)^44^. Although highly context dependent, a few forms of ICD include immunogenic apoptosis, necroptosis, and pyroptosis^45–47^. While the exact mechanisms may vary between different forms of ICD, they all must result in the stimulation of innate immune cells via DAMPs, as well as the successful processing and presentation of tumor-associated antigens to cytotoxic T lymphocytes by antigen presenting cells^48–51^. Having this dual effect on innate and adaptive immune compartments allows immunologically “cold” tumors—characterized by poor immune infiltration—to be recognized and targeted by the host immune system^52^.

As noted above, metabolic deregulation is a hallmark of PDAC cells, which meet their biosynthetic demands through metabolic rewiring and the activation of varied nutrient acquisition programs. While these unique metabolic adaptations represent mechanisms of PDAC tumor cell survival, they may also present therapeutic targets to trigger ICD^53–59^. This approach capitalizes on PDAC’s inherent metabolic dependencies by targeting malignant cells while simultaneously overcoming microenvironmental immunosuppression. Based on these considerations, we set forth to investigate metabolism-induced ICD.

Using a diverse panel of syngeneic murine cancer cell lines, we first investigated whether features of ICD could be uniquely elicited by perturbation of amino acid metabolic pathways. Among the amino acids analyzed, restriction of cystine induced canonical features of ICD in multiple cancer cell types, including PDAC. We subsequently demonstrated that the ICD features of cysteine deprivation were a consequence of ferroptosis, a metabolic form of cell death marked by deregulated management of lipid oxidation. Using lipidomic and metabolomic analyses of ferroptotic PDAC cells, we observed the selective release of immunomodulating metabolites and lipids. Despite the clear induction of ICD features in vitro, our in vivo tumor vaccination studies illustrated that ferroptotic cell death inversely contributes to the creation of a tumor-protective environment. This was evidenced by tumor outgrowth at vaccination sites, myeloid cell infiltration, and TME alterations. The sum of our work adds to the body of evidence revealing the complex nature of ferroptosis in tumor immunity and emphasizes the influence of cell models and context in studying cell death and immunity^60–63^.

## Methods

### Cancer cell lines and cell culture

Cells were cultured in a humidified incubator at 37°C with 5% CO2. Cell propagation media for all cell lines involved in this study consisted of RPMI 1640 (Gibco, 11965092) with 10% FBS (Corning, 35-010-CV) unless otherwise indicated. PBS (Gibco, 10010023) was used for all cell washing steps unless noted otherwise. All cell lines were routinely screened for mycoplasma using MycoAlert PLUS Mycoplasma Detection Kit (LT07-710). All data included in this manuscript were generated with mycoplasma negative cell cultures. KPC7940b murine cells were obtained under a material transfer agreement by G. Beatty. The murine EG7 cell line used herein was purchased from the American Type Culture Collection (ATCC). CT26 and GL261 lines were a gift from Y. Shah. The identity of all cell lines were confirmed by STR profiling.

### Flow Cytometry

#### Viability and exposure of ecto-calreticulin

Cells were washed once in ice-cold Annexin Binding Buffer (ThermoFisher, V13246), then resuspended in ice-cold staining solutions containing Annexin V buffer plus Calreticulin-FITC (Novus Biologicals, NBP1-47518F) at a 1:100 dilution or Annexin V-Pacific Blue (ThermoFisher, A35122) at a 1:300 dilution. After 40 minutes of incubation, staining solutions were removed and cells washed in ice cold Annexin Binding Buffer. Next, cells were resuspended in ice-cold Annexin V binding buffer containing a 7-AAD viability staining solution (ThermoFisher, 00-6993-50), followed by flow cytometry analysis (Bio-Rad Ze5 Cell Sorter).

#### Lipid ROS measured by C-11 BODIPY

Cell were stained in prewarmed phenol-red free media with low FBS (0.2% FBS) using 2 μM C-11 BODIPY (ThermoFisher, D3861) for 30 minutes 37°C and 5% CO_2_. Cells were then washed in ice-cold PBS twice before detachment via trypsin. After neutralization and removal of trypsin, staining with 1 μM Sytox Blue (ThermoFisher, S34857) was performed. Cells were promptly analyzed using a Bio-Rad Ze5 Cell Sorter.

#### Extracellular ATP

Cells were plated and treated according to assay specific conditions. At endpoint, extracellular ATP was measured using ENLITEN ATP Assay System Bioluminescence Detection Kit (Promega, FF2000) according to the manufacturer’s protocol.

### Amino acid dropout studies

#### Assays

Media conditions for assays with EG7 cells included either all the amino acids in standard RPMI (indicated below), or media with omission of arginine, cystine, glutamine, leucine, serine (+glycine), or tryptophan. Media conditions for assays with CT26 and GL261 cells included either all the amino acids in standard RPMI, or media with omission of cystine, glutamine, or serine (+glycine). EG7 cells lines were washed and seeded in 96 well plates at 40,000 cells per well in standard RMPI or amino acid dropout media conditions, then assessed for viability, calreticulin, and extracellular ATP release. GL261 and CT26 cells were seeded in six-well plates at 150,000 and 120,000 respectively and allowed to adhere overnight. The following day, GL261 and CT26 cells were washed and their media replaced with either standard RMPI or amino acid dropout media conditions. After 24 hours of incubation, cells were assessed for viability, calreticulin, and extracellular ATP release.

#### Amino acid dropout media

These assays used standard RPMI with addition of 10% FBS (Corning, 35-010-CV) for conditions including all amino acids, or an in-house, RPMI media formulation that matched standard RPMI in composition and concentration of all components minus the single amino acids indicated in each study. The following amino acids were purchased from Sigma-Aldrich and were used at the indicated concentrations: L-arginine, 200 mg/mL (A8094-25G); L-Asparagine (anhydrous), 50 mg/mL (A4159-25G); L-Aspartic Acid, 20 mg/mL (A7219-100G); L-Cystine dihydrochloride, 65.2 mg/mL (C6727-25G); L-Glutamine, 300 mg/mL (G8540-25G); L-Glutamic Acid, 20 mg/mL (G8415-100G); Glycine, 10 mg/mL (G8790-100G); L-Histidine, 15 mg/mL (H6034-25G); Hydroxy-L-Proline, 20 mg/mL (H5534-10MG); L-Isoleucine, 50 mg/mL (I7403-25G); L-Leucine 50, mg/mL (L8912-25G); L-Lysine•HCl, 40 mg/mL (L8662-25G); L-Methionine, 15 mg/mL (M5308-25G); L-Phenylalanine, 15 mg/mL (P5482-25G); L-Proline, 20 mg/mL (P5607-25G); L-Serine, 30 mg/mL (S4311-25G); L-Threonine, 20 mg/mL (T8441-25G); L-Tryptophan, 5 mg/mL (T8941-25G); L-Tyrosine +2Na +2H2O, 28.83 mg/mL (T1145-25G); L-Valine, 20 mg/mL (V0513-25G).

#### Isolation of BMDCs

BMDCs were derived from C57BL/6J mice purchased from Jackson Laboratory. Cells were flushed with PBS from the isolated femurs and tibias of 6-8 week-old mice and cultured for seven days in a BMDC media formulation containing the following: RPMI-1640 media (Gibco, 11875135), 10% FBS (Corning, 35-010-CV), penicillin-streptomycin 100 U/mL (Gibco, 15140122), 2 mM L-glutamine (Gibco, 25030081), 50 μM 2-mercaptoethanol (ThermoFisher, 31350010), 1X non-essential amino acids (Gibco, 11140076), 1 mM sodium pyruvate (Gibco, 11360070), and 20 ng/mL GM-CSF (Abcam, ab9742). Every two days cultures were supplemented with fresh media. After seven days of culture cells were collected and used for experiments. BMDCs are identified as floating and loosely attached cells on the dish following culture of bone marrow cells.

#### BMDC phagocytosis and maturation assays

Prior to coculture for phagocytosis assays, EG7 cells were stained with 1μM Cell Tracker Green CMFDA Dye (Invitrogen, C2925) in phenol-free, serum-free RPMI-1640 media and cultured for 48 hours in respective media conditions at 500,000 per well in 24-well plates. For coculture, BMDCs were plated at 200,000 cells per well in 6-well plates in the aforementioned BMDC media. EG7 cells were then assayed for viability by trypan blue staining (Gibco, 1525006) and added at indicated ratios with BMDCs for 24 hours before harvest and subsequent flowcytometry. For BMDC maturation assays, EG7 cells were plated at 500,000 cells per well in a 24-well plate according to assay specific media conditions. After 48 hours, BMDCs were plated at 200,000 cells per well in 6-well plates in the aforementioned BMDC media. Target cells were assayed for viability and added at indicated ratios with BMDCs for 24 hours before harvest and subsequent flowcytometry.

BMDCs assessed in phagocytosis assays were stained for 35 minutes in cold PBS with APC-CD11c (BD Biosciences, 55026). After one wash, BMDCs were resuspend cells in 1μM Sytox blue (ThermoFisher, S11348). Analysis was performed using FlowJo (v.10.0.8) software. BMDCs were sorted using a Bio-Rad Ze5 Cell Sorter. Gating strategy: Single cells were obtained by gating on FSC-A versus FSC-H, followed by SSC-A versus SSC-H. Dead cells were excluded by SYTOX Blue. BMDCs that successfully phagocytized target cells were identified as CD11c+CMFDA+. BMDCs assessed in maturation assays were stained for 35 minutes in cold PBS with APC-CD11c (BD Biosciences, 550261), PeCy7-CD86 (BD Biosciences, 560582), PE-Cy7-anti-CD80 (eBioscience, 25080182), and FITC-MHCII (eBioscience,11532181). After one wash, BMDCs were resuspend cells in 1μM Sytox blue (ThermoFisher, S11348). Analysis was performed using FlowJo (v.10.0.8) software. BMDCs were sorted using a Bio-Rad Ze5 Cell Sorter. Gating strategy: Single cells were obtained by gating on FSC-A versus FSC-H, followed by SSC-A versus SSC-H. Dead cells were excluded by SYTOX Blue. Mature BMDCs cells were identified as a percent of CD11c+ CD86+ MHCII+ and CD11c+ CD80+ MHCII+.

#### Cell death-inducing reagents

Erastin (Cayman, 17754), RSL3 (Cayman, 19288), IKE (Cayman, 27088), Ferrostatin-1 (Cayman, 17729), Bafilomycin A1 (Cayman, 11038), Necrosulfonamide (Sapphire North America, 20844), Z-VAD(Ome)-FMK (Cayman,14463), Necrostatin-1 (Cayman, 11658), Mitoxantrone (Sapphire North America, 14842).

### EG7 cell assays

#### Cystine viability assay

Media conditions were made fresh before running assays. Cystine dropout media was formulated in house using RPMI 1640 Medium Modified (Sigma, R75130100ML) with add back of L-Methionine (Sigma-Aldrich, M5308-25G) at 15 mg/mL, L-Glutamine (G8540-25G) at 300 mg/mL, and L-Cystine (Sigma-Aldrich, 57579-5ML-F) at 65.2 mg/L, 6.52 mg/L, 0.652 mg/L, 0.0652 mg/L, or 0.00652 mg/mL. All media was supplemented with 10% FBS. Cells were washed and seeded in 96-well plates at 40,000 cells per well in 200 µL of each media condition with or without 2 µM Fer-1 for 96 hours. At end point, cell viability was measured using CellTiter-Glo® 2.0 Cell Viability Assay.

#### RSL3 and Erastin viability, ecto-calreticulin, and extracellular ATP assays

For viability and ecto-calreticulin measurements, cells were washed and seeded in 96-well plates at 40,000 cells per well in 200 µL of media containing each indicated drug condition with or without 2 µM Fer-1 for 24 hours. At end point, cells were centrifuged and 100 µL of media was taken for extracellular ATP measurements. Cells were then processed for flow cytometry to measure ecto-calreticulin or cell viability.

#### Lipid peroxidation assay

Minus cystine media was formulated in house using RPMI 1640 Medium Modified (Sigma, R75130100ML) with add back of L-Methionine (Sigma-Aldrich, M5308-25G) at 15 mg/mL, L-Glutamine (G8540-25G) at 300 mg/mL. Plus cystine media conditions consisted of the same media as the minus cystine condition along with add back of L-Cystine (Sigma-Aldrich, 57579-5ML-F) at 65.2 mg/L. The 3 µM Erastin media condition consisted of the same media as the plus cystine condition. All media was supplemented with 10% FBS. Cells were washed and seeded in 96-well plates at 40,000 cells per well in 200 µL of each media condition with or without 2 µM Fer-1 for 12 hours. At end point cells were processed for flow cytometry to measure C-11 BODIPY.

#### Generation of GPX4 knockout cells

GPX4 CRISPR–Cas9 constructs were generated using the expression vector pSpCas9(BB)-2A-Puro (PX459) V2.0, which was obtained from Addgene (Plasmid #62988). As described formerly^64^, the restriction enzyme BbsI was used to cut the plasmid followed by ligation of mouse glutathione peroxidase 4 sgRNA sequences. Oligonucleotide pairs were obtained from the Human GeCKO Library (v2, 3/9/2015). The day prior to transfection, KPC7940b cells were seeded at 200,000 cells per well in a 6-well plate. Lipofectamine® LTX Reagent (Invitrogen, 15338100) in conjunction with PLUS™ (Invitrogen, 11514015) was used to transfect cells with 2.5 μg of pSpCas9-GPX4 according to the manufacturer’s instructions. Selection with 2 μg mL-1 puromycin began 24 hours later. Selection was continued with replacement of 2 μg mL-1 puromycin media every two days until death of all non-transfected cells. Single cell clones were selected and expanded from successfully transfected cell lines. At the time of transfection and during all subsequent culture of GPX4 KO cells, media was supplemented with 2 μM Ferrostatin-1.

### KPC7940b cell assays

#### Time course of extracellular ATP and CXCL1

KPC7940b wild type and GPX4 knockout cells were seeded at 9×10^5^ cells in 10 cm plates in 10 mL DMEM (Gibco, 11965092) medium with 10% FBS (Corning, 35-010-CV). 18 hours later, cells were washed, and media was replaced with 6 mL media and the indicated conditions. Samples of supernatants from each plate were taken at 0, 1-, 2-, 4- and 8-hour time points, centrifuged at 300 g for 5 minutes at 4°C and then promptly divided for measuring either extracellular ATP, or CXCL1 by ELISA. Extracellular ATP was measured by ENLITEN ATP Assay System Bioluminescence Detection Kit (Promega, FF2000) according to the manufacturer’s protocol.

#### Analysis of calreticulin exposure and lipid peroxidation

KPC7940b wild type and GPX4 knockout cell lines were seeded at 2×10^5^ cells per well in a 6-well plate. 24 hours later, cells were washed twice with 3 mL PBS. After washing, cells were treated with 3 mL of media containing their respective conditions. After four hours, cells were promptly collected and assessed for ecto-calreticulin exposure and lipid peroxidation by flow cytometry.

#### Analysis of viability

KPC7940b wild type and GPX4 knockout cell lines were washed and seeded in 96-well plates at 2×10^4^ cells per well in 200 µL of their indicated media condition with or without 2 µM Fer-1 for 72 hours. At end point, cell viability was measured using CellTiter-Glo® 2.0 Cell Viability Assay.

#### Cell death inhibitor assays

KPC7940b wild type cells were washed and seeded in 96-well plates at 4×10^4^ cells per well in 200 µL of media. Cells were co-treated with the indicated cell death inhibitors and either 5 µM imidazole ketone erastin, 1 µM RSL3, or DMSO vehicle. 48 hours later, at end point, cell viability was measured using CellTiter-Glo® 2.0 Cell Viability Assay.

#### Colony Forming Assay

KPC7940b wild type and GPX4 knockout cell lines were washed and seeded in 6-well plates at 250 cells per well in 3 mL of media with their indicated treatment conditions. After 6 days, media was removed, cells were washed in ice-cold PBS and fixed with 100% methanol for 10 minutes. After removal of methanol, cells were stained with 0.5% crystal violet solution for 15 minutes.

Crystal violet was then aspirated, followed by repeated washing steps to remove the remaining crystal violet.

#### Dose response curves

KPC7940b cells were washed and seeded in 96-well plates at 4×10^4^ cells per well in 200 µL of media with either imidazole ketone erastin, RSL3, or cystine media conditions, each with or without 2 µM Fer-1. At endpoint, 48 hours later, cell viability was measured using CellTiter-Glo® 2.0 Cell Viability Assay. Cystine dropout media was formulated in house using RPMI 1640 Medium Modified (Sigma, R75130100ML) with add back of L-Methionine (Sigma-Aldrich, M5308-25G) at 15 mg/mL and L-Glutamine (G8540-25G) at 300 mg/mL. L-Cystine (Sigma-Aldrich, 57579-5ML-F) was added back at various concentrations, where 100% cystine is equal to 65.2 mg/L. All media was supplemented with 10% FBS. IC_50_ values were calculated using Prism analyzation tools.

### Metabolomic profiling

#### Metabolomics sample preparation

For both media supernatant and cell pellet metabolomic analysis, PDAC cells were plated in triplicate in 10 cm plates at a density of 9×10^5^ cells in DMEM (Gibco, 11965092) medium with 10% FBS (Corning, 35-010-CV). Two additional plates were prepared in parallel for protein quantification, and subsequent normalization of samples, as well as for flow cytometry to measure viability by 7-AAD. After incubating overnight, the culture medium was removed, cells were washed with PBS, and fresh medium was added containing the indicated conditions. Cells were then incubated for four hours, at which point supernatants and pellets were collected for metabolite extraction. To collect metabolites from media supernatants, 1.5 mL of culture medium was taken from each plate, transferred to a tube, and spun at 300 g for 5 minutes at 4°C. Next, 200 µL of supernatant was transferred to a new tube, followed by the addition of 800 µL of ice-cold methanol. For metabolite extraction from cell pellets, the remaining medium was aspirated, and the cells were washed once with 4 mL PBS. Next, 4 mL of ice-cold methanol/water 1:4 (v/v) was added, and the plate was incubated on dry ice for 10 minutes. The resulting cell lysates were then transferred into tubes for centrifugation at 12,000 g. For each experimental group, the volume of supernatant collected from cell lysates for drying was adjusted according to the protein content measured from the cell pellets of the parallel plate. Samples were dried using a SpeedVac Vacuum Concentrator (model: SPD1030).

#### Targeted Metabolomics

The dried supernatants were reconstituted in methanol diluted in water (1:1) and analyzed by LC-MS according to a previously described protocol^65^. Data processing was performed using Agilent MassHunter Workstation Quantitative Analysis for QQQ software (version 10.1, build 10.1.733.0). Heatmaps were generated using MetaboAnalyst 6.0^66^.

Raw data are provided as **Supplementary File 1**.

### Lipidomic profiling

#### Lipidomics sample preparation

For both media supernatant and cell pellet lipidomic analysis, PDAC cells were plated in triplicate in 15 cm plates at a density of 1×10^7^ cells in DMEM (Gibco, 11965092) medium with 10% FBS (Corning, 35-010-CV). An additional plate was prepared in parallel for protein quantification and subsequent normalization of samples. After incubating overnight, the culture medium was removed, cells were washed with PBS, and fresh medium was added containing the indicated conditions. Cells were then incubated for four hours, at which point supernatants and pellets were collected for lipid extraction. To collect lipids from media supernatants, 1.5 mL of culture medium was taken from each plate, transferred to a tube, and centrifuged at 300 g for 5 minutes at 4°C. After centrifugation, 1 mL of supernatant from each replicate were immediately transferred to a new tube and frozen at −80°C along with 1 mL of cell-free culture media as a control. To collect lipids from cell pellets, the remaining medium was aspirated, and the cells were washed twice with 15 mL PBS. Cells were collected from plates using trypsin, followed by centrifugation at 300 g for 5 minutes at 4°C. Trypsin was then removed and cell pellets were washed twice in 15 mL PBS. After aspirating the second wash, cells pellets were resuspended in 1 mL PBS containing 200 μM diethylenetriaminepentaacetic acid (Sigma, D6510-10G). Next, one part methanol was added, after which samples were promptly frozen at −80°C. Protein content was measured using the cell pellets of the parallel plate. These protein measurements were used for normalizing lipids from cell pellets. Samples were transported to Cayman Chemical on dry ice.

#### Targeted and untargeted lipidomics

Lipidomics analysis was performed as detailed below by Cayman Chemical Company, Ann Arbor, Michigan. Upon receipt, samples were stored at −80°C until the time of analysis. After thawing, lipids were extracted using a methyl-tert-butyl ether (MTBE)-based liquid-liquid extraction method. Samples were thawed on ice in the original tubes before adding 500 μL PBS/methanol 1:1 (v/v) and then 100 μL methanol containing 50 ng each of the following internal standards for targeted lipidomics: PC(15:0/18:1-d7), PC(15:0/18:1-d7), PE(15:0/18:1-d7), PE(15:0/18:1-d7), PG(15:0/18:1-d7), PG(15:0/18:1-d7), PI(15:0/18:1-d7), PI(15:0/18:1-d7), PS(15:0/18:1-d7), PS(15:0/18:1-d7), PC(15:0/18:1-d7), PE(15:0/18:1-d7), PG(15:0/18:1-d7), PI(15:0/18:1-d7), and PS(15:0/18:1-d7). For untargeted metabolomics, 500 μL PBS/methanol 1:1 (v/v) and then 100 μL methanol containing 50 ng each of the following internal standards was added: TG(15:0/18:1-d7/15:0), DG(15:0/18:1-d7/0:0), Cer(d18:1-d7/15:0), SM(d18:1/18:1-d9), PG(15:0/18:1-d7), PC(15:0/18:1-d7), PI(15:0/18:1-d7), PS(15:0/18:1-d7), LysoPC(18:1-d7), LysoPE(18:1-d7), PE (15:0/18:1-d7). Samples were then transferred into 8-mL screw-cap tubes, and then 1.125 methanol and 5 mL MTBE were added. After vigorous mixing, samples were incubated at room temperature on a tabletop shaker for 45 min. For phase separation, 1.25 mL water was added, and samples were vortexed and centrifuged for 15 min at 2000 x g. The upper organic phase of each sample was carefully removed using a Pasteur pipette, transferred into an empty glass round-bottom tube, and dried under vacuum in a SpeedVac concentrator.

The dried lipid extracts were resuspended in 200 μL HPLC mobile phase A/mobile phase B 3:1 (v/v) for targeted LC-MS/MS analysis. For untargeted LC-MS/MS analysis, resuspended extracts were dried again under vacuum in a SpeedVac concentrator, then resuspended in 100 μL 1-butanol/methanol 1:1 (v/v). For targeted LC-MS/MS, extracted lipid samples (20 µL) were injected onto a Solex ExionLC Integrated System coupled to a Sciex QTrap 6500+ mass spectrometer. Lipid separation was performed using a reverse-phase Kinetex HILIC core shell column (2.6 µm, 150 × 2.1 mm, Phenomenex) maintained at room temperature, with a flow rate of 200 µL/min. The mobile phases consisted of hexane/isopropanol (30:40, v/v) (Mobile Phase A) and hexane/isopropanol/water (30:40:7.5, v/v/v) with ammonium acetate (Mobile Phase B). The chromatographic gradient was as follows: 0 min, 25% B; 1 min, 25% B; 4 min, 60% B; 7 min, 85% B; 10 min, 95% B; 11 min, 25% B; 15 min, 25% B for re-equilibration. Mass spectrometric analysis was performed using electrospray ionization (ESI) in both positive and negative ion modes, with polarity switching. The electrospray voltage was set to 4500 V for both positive and negative modes, with a source temperature of 450°C. Ion source gas 1 and gas 2 flow rates were set to 35 psi each, and the curtain gas was set to 35 psi. The collision gas was set to low, with collision energy set to −30 eV (negative mode) or +45 eV (positive mode). The declustering potential was set to ±100 V, entrance potential to ±10 V, and collision cell exit potential to −7.5 V (negative mode) or +10 V (positive mode). Data were acquired in MRM (Multiple Reaction Monitoring) mode with a dwell time of 25 ms. For untargeted LC-MS/MS, extracted lipid samples (5 µL) were injected onto an Ultimate 3000 UPLC system (Thermo Scientific) coupled to a Q-Exactive Plus Orbitrap mass spectrometer (Thermo Scientific). Lipid separation was performed using a reverse-phase Accucore C30 column (2.6 µm, 150 x 2.1 mm, Thermo Scientific) maintained at 40 °C, with a flow rate of 300 µL/min. The mobile phases consisted of acetonitrile/water/formic acid (60:40:0.1, v/v/v) with 10 mM ammonium formate (Mobile Phase A) and acetonitrile/isopropanol/formic acid (10:90:0.1, v/v/v) with 10 mM ammonium formate (Mobile Phase B). The chromatographic gradient was as follows: 0 min, 30% B; 5 min, 43% B; 5.1 min, 50% B; 14 min, 70% B; 14.1 min, 70% B; 21 min, 90% B; 28 min, 98% B; 33 min, 30% B for re-equilibration. Mass spectrometric analysis was performed using electrospray ionization (ESI) in both positive and negative ion modes, with polarity switching. The spray voltage was set to 3.0 kV for positive mode and 3.2 kV for negative mode, with a capillary temperature of 350 °C. Sheath gas and auxiliary gas flow rates were set to 60 and 20 (arbitrary units), respectively, and the S-lens RF level was 45. The collision energy was set to 28 eV. Data were acquired in both full MS and data-dependent MS² (dd-MS²) modes, with a mass resolution of 70,000 (MS) and 35,000 (dd-MS²). The scan range was 200–2000 m/z, with an AGC target of 1e6 (200 ms, MS; 1e6 (300 ms), dd-MS²), TopN set to 8, and an isolation window of 1.0 m/z.

Data processing was performed by Cayman Chemical Company using Lipostar software (v. 1.3.1b18; Molecular Discovery) for feature detection, noise and artifact reduction, alignment, normalization and lipid identification. Heatmaps were generated using MetaboAnalyst 6.0^66^.

Raw data are provided as **Supplementary Files 2,3**.

#### Western blotting

Cell lysates were collected using RIPA buffer (Sigma-Aldrich, R0278) with protease and phosphatase inhibitors on ice. Lysate protein concentrations were quantified by bicinchoninic acid assays (ThermoFisher, 23227). Lysates were diluted in loading dye (ThermoFisher, NP0007), reducing agent (ThermoFisher, NP0009), and RIPA buffer to a protein concentration of 20 ug per sample prior to loading on a NuPAGE 4–12% Bis-Tris gel (ThermoFisher, NP0336BOX) along with SeeBlue Plus2 protein ladder (Invitrogen, LC5925) for electrophoresis at 150 V for 45 minutes in NuPAGE MOPS SDS Running Buffer (ThermoFisher, NP0001).

Protein was then blotted to a methanol activated PVDF membrane (Millipore, IPFL00010) in transfer buffer (Invitrogen, NP00061) at 25 V for one hour. Membranes were incubated at room temperature in 5% nonfat milk blocking buffer (Bio-Rad, 1706404) for one hour before incubation in the indicated antibody overnight at 4’ C. Membranes were washed three times in tris-buffered saline (Bio-Rad, 1706435) with 0.1% Tween-20 (Sigma-Aldrich, 9005-64-5) for 5 minutes per wash in between primary, secondary and ECL Reagent (Bio-Rad, 705062) incubations. Images were captured with a Bio-Rad ChemiDoc Imaging System (Image Lab Touch Software version 2.4.0.03). The following primary and secondary antibodies were used: anti-GPX4 (Abcam, ab125066), anti-Vinculin (Cell Signaling, 13901S), both at a 1:1,000 dilution, and anti-rabbit-HRP (Cell Signaling, 7074S) at a 1:10,000 dilution.

Raw data are provided as **Supplementary File 4**.

#### T-Cell Assay

KPC7940b wild type and GPX4 knockout cells were plated at 9×10^5^ cells per well in 10 cm plates with 10 mL of RPMI 1640 (Gibco, 11965092) with 10% heat-inactivated FBS (Corning, 35-010-CV). 18 hours later cells were washed twice with 10 mL PBS followed by replacement with 6 mL of the indicated media conditions. Cells were left to incubate for four hours, after which the conditioned media was collected, centrifuged, and filtered through a 0.45 μm filter. The following supplements were next added to the conditioned media for culturing of CD8^+^ T cells: 1% penicillin/streptomycin (Gibco, 15140163), 50 μM 2-mercaptoethanol (Gibco, 21985023), 2 mM L-Glutamine (Gibco, 25030081), 10 mM HEPES (Gibco, 15630080), and 200 U/mL IL-2 (PeproTech, 212-12-20UG). Naïve CD8 T cells were isolated from mouse spleens and lymph nodes by magnetic bead separation (Miltenyi Biotec, 130-096-543) following the manufacturers’ protocols. CD8^+^ T cells were then seeded in a 96-well plate that was coated in a PBS solution of 1 μg/mL aCD3 (Biolegend, 100340) and 5 μg/mL aCD28 (Biolegend, 102116). T cells were then incubated for 48 hours in 100 μL of their respective media, after which they were analyzed for proliferation using the CellTrace Violet Cell Proliferation Kit (Invitrogen, C34557) and activation (Millipore Sigma, MABF1556) by flow cytometry using a Bio-Rad Ze5 Cell Sorter. Data were analyzed with FlowJo v11 Software.

#### Mice

Animal experiments were approved by the University of Michigan Institutional Animal Care and Use Committee following PRO00010606. Both C57BL/6 (strain 000664, aged 6-8 weeks) and NSG mice (strain 005557, aged 6-8 weeks) were obtained from Jackson Laboratory. Mice were housed in a pathogen-free animal facility with a 12-hour light/12-hour dark cycle, 30-70% humidity, and temperatures 20-23 °C. Mice were housed at no more than five per cage. Sample size calculations were performed in advance.

#### Prophylactic vaccination study

In each arm of this study, 10 C57BL/6J mice were used. EG7 cells were treated with either 10 μM RSL3, 10 μM Mitoxantrone, or Oxaliplatin 10 μM for 24 hours, after which RSL3 and Oxaliplatin treated cells were assessed by 7-AAD for viability and Mitoxantrone treated cells were assessed by Trypan Blue for viability. Mice were vaccinated with a total of 1×10^6^ cells per treatment condition, where RSL3 treated cells were 24% viable, Mitoxantrone treated cells were 29% viable, and Oxaliplatin treated cells were 28% viable. Nine days after mice received a vaccination on their left flank, mice received a subcutaneous tumor challenge of 6×10^5^ EG7 cells on their right flank. Subcutaneous tumor injections consisted of a 100 μL cell suspension in serum-free DMEM/Matrigel (Corning, 354234) 1:1 (v/v). Tumor measurements were taken starting 10 days post tumor challenge. For this study, all mice were taken down on day 23 post tumor challenge, unless tumor ulcerations were observed or tumor volumes exceeded 2.0×10^3^ mm^3^, at which point mice were euthanized.

### Tumor vaccine studies

#### KPC7940b GPX4 knockout cell line study and immunohistochemistry

For both arms of this study, five C57BL/6J mice were used. Four hours prior to their preparation for subcutaneous tumor injections, KPC7940b GPX4 knockout cells were withdrawn from their standard culture conditions of DMEM supplemented with 2 μM ferrostatin-1 and 10% FBS and grown in standard DMEM with 10% FBS for four hours. The purpose of this step was to induce ferroptotic cell death as performed in other experiments herein. Subcutaneous tumor injections contained a total of 5×10^5^ KPC7940b GPX4 knockout cells along with an equivalent number of KPC7940b wild type cells as a control. Subcutaneous tumor injections consisted of a 100 μL cell suspension in serum-free DMEM/Matrigel (Corning, 354234) 1:1 (v/v). Tumor measurements were taken starting three days after setting subcutaneous tumors. At endpoint, tumors were fixed in 10% neutral buffered formalin for 48 hours, then embedded in paraffin to create formalin-fixed paraffin-embedded FFPE blocks. Serial sections, each 4 μm thick, were cut from the FFPE blocks. Sections were deparaffinized in Histoclear and rehydrated through a graded ethanol series to distilled water. Slides were stained in hematoxylin, rinsed in tap water, and differentiated in acid alcohol if necessary. Following bluing in tap water, sections were counterstained in eosin, dehydrated through graded alcohols, cleared in Histoclear, and mounted with coverslips using Toluene. For immunostaining, sections were subjected to antigen retrieval using citrate buffer (pH 6.0) or Tris-EDTA buffer (pH 9.0) at 95-100°C, followed by cooling to room temperature. Endogenous peroxidase activity was quenched with 3% hydrogen peroxide. After blocking with 5% normal serum, sections were incubated overnight at 4°C with primary antibodies against GPX4, CK19, CD3 (eBioscience, 100340), F4/80 (eBioscience, 15480182), or αSMA (Millipore-Sigma, A5228) at manufacturer-recommended dilutions. The following day, slides were washed and incubated with appropriate biotinylated secondary antibodies for 30 minutes at room temperature. Signal was visualized using a DAB substrate kit, and sections were counterstained with hematoxylin, dehydrated, cleared, and mounted.

#### KPC7940b cell line study in BL/6 mice and CyTOF analysis

In this study, 10 C57BL/6 mice were used in the control and RSL3 arms, and seven mice were used in the Mitoxantrone arm. KPC7940b cells were treated with either vehicle (control), 10 μM RSL3, or 10 μM Mitoxantrone for 24 hours, after which cells were assessed by Trypan Blue for viability. Subcutaneous injections were prepared such that RSL3 treated and Mitoxantrone arms received a total of 1×10^6^ cells, where RSL3 treated cells were 27% viable and Mitoxantrone treated cells were 29% viable. Injections for the control arm were prepared to have 2.5×105 viable cells. Subcutaneous tumor injections consisted of a 100 μL cell suspension in serum-free DMEM/Matrigel (Corning, 354234) 1:1 (v/v). Tumor measurements were taken starting 9 days after setting subcutaneous tumors. All mice were taken down 24 days post tumor challenge. At endpoint, three tumors from each condition were analyzed by Cytometry by Time-of-Flight analysis. The processing of tissues and data analysis closely followed protocols that were previously described in detail^67^. The following antibodies from Standard BioTools were used at listed dilutions: CD45, 1:200 (3089005C); CD11b, 1:300 (3148003B); CD206, 1:200 (3169021B); CD3, 1:100 (3152004B); CD8, 1:200 (3168003B); CD11c, 1:100 (3209005B); Ly6c, 1:400 (3150010B); Ly-6G, 1:400 (3141008B); iNOS, 1:100 (3161011B); Arg1, 1:100 (3164027D); F4/80, 1:100 (3146008B); CD4, 1:200 (3145002B).

#### KPC7940b cell line study in BL/6 vs. NSG mice and flow cytometry analysis

For this study, three conditions—Vehicle (control), RSL3, or freeze/thaw—were tested both immunocompetent C57BL/6 and immune incompetent NOD *scid* gamma (NSG) mice. For the C57BL/6 arm, there were 10 mice per condition, and in the NSG arm there were five mice per condition. Subcutaneous tumor injections in the freeze/thaw condition consisted of KPC7940b cells subject to two freeze/thaw cycles. For the other two conditions, KPC7940b cells were treated with either vehicle (control) or 10 μM RSL3 for 24 hours. Subcutaneous tumor injections were prepared such that RSL3 treated, and freeze/thaw arms received a total of 1×10^6^ cells, where RSL3 treated cells were approximately 30% viable and freeze/thaw treated cells were approximately 30% viable as determined by Trypan Blue. Tumor injections for the control arm were prepared to have 3×10^5^ viable cells. Subcutaneous tumor injections consisted of a 100 μL cell suspension in serum-free DMEM/Matrigel (Corning, 354234) 1:1 (v/v). Tumor measurements were taken starting eight days after setting subcutaneous tumors. All mice were taken down 24 days post tumor challenge. At end point, tumors from each condition were analyzed by flow cytometry. Tumors mechanically dissociated using scissors in sterile PBS, followed by centrifugation and resuspension in 1 mg/mL Collagenase V (Sigma) for enzymatic digestion at 37°C for 30 minutes. Digestion was quenched by adding DMEM supplemented with 10% FBS. After sequentially filtering the cell suspension through 500 μm, 100 μm, and 40 μm strainers to obtain a single-cell suspension. Red blood cells were lysed, and the remaining cells were washed with PBS. Cells were then blocked and stained with antibodies in FBS. After staining, cells were washed in FACS buffer (1% Bovine Serum Albumin and 1 mM ethylenediaminetetraacetic acid in PBS) and analyzed on a MoFlo Astrios flow cytometer (Beckman Coulter). The following antibodies were used: MHCII (eBioscience,11532181), CD45.1 (eBioscience, 11045381), CD8a (BD Biosciences, 557654), CD11b (BD Biosciences, 557657), F4/80 (eBioscience, 15480182), CD11c (eBioscience, 562782), TCRb (eBioscience, 12596182).

#### Statistical analysis

Data are presented as the mean ± SD from technical or biological replicates as indicated in legends. Statistical analyses were performed using GraphPad Prism 10.5.0 (GraphPad Software Inc.). Differences between experimental and control groups were assessed for significance using unpaired t-tests with Welch’s correction. Where comparisons are made among multiple experimental groups and controls, one-way analysis of variance (ANOVA) was used. A p-value below 0.05 was considered statistically significant.

## Results

### Metabolic screen identified that cystine dropout induces features of immunogenic cell death

The objective of this study was to identify metabolic mechanisms that regulate ICD in cancer. We elected a panel of syngeneic murine tumor cell lines representing diverse cancer types. Syngeneic cell lines are genetically identical to inbred mouse strains and can be transplanted into immune competent mice without rejection. This enables the study of immune-tumor interactions *in vivo*. With this panel, we analyzed the role of amino acids, based on their druggable, diet-amendable, and experimentally tractable features, and how amino acid dropout impacted features of ICD (**Fig. 1A**).

**Figure 1.**
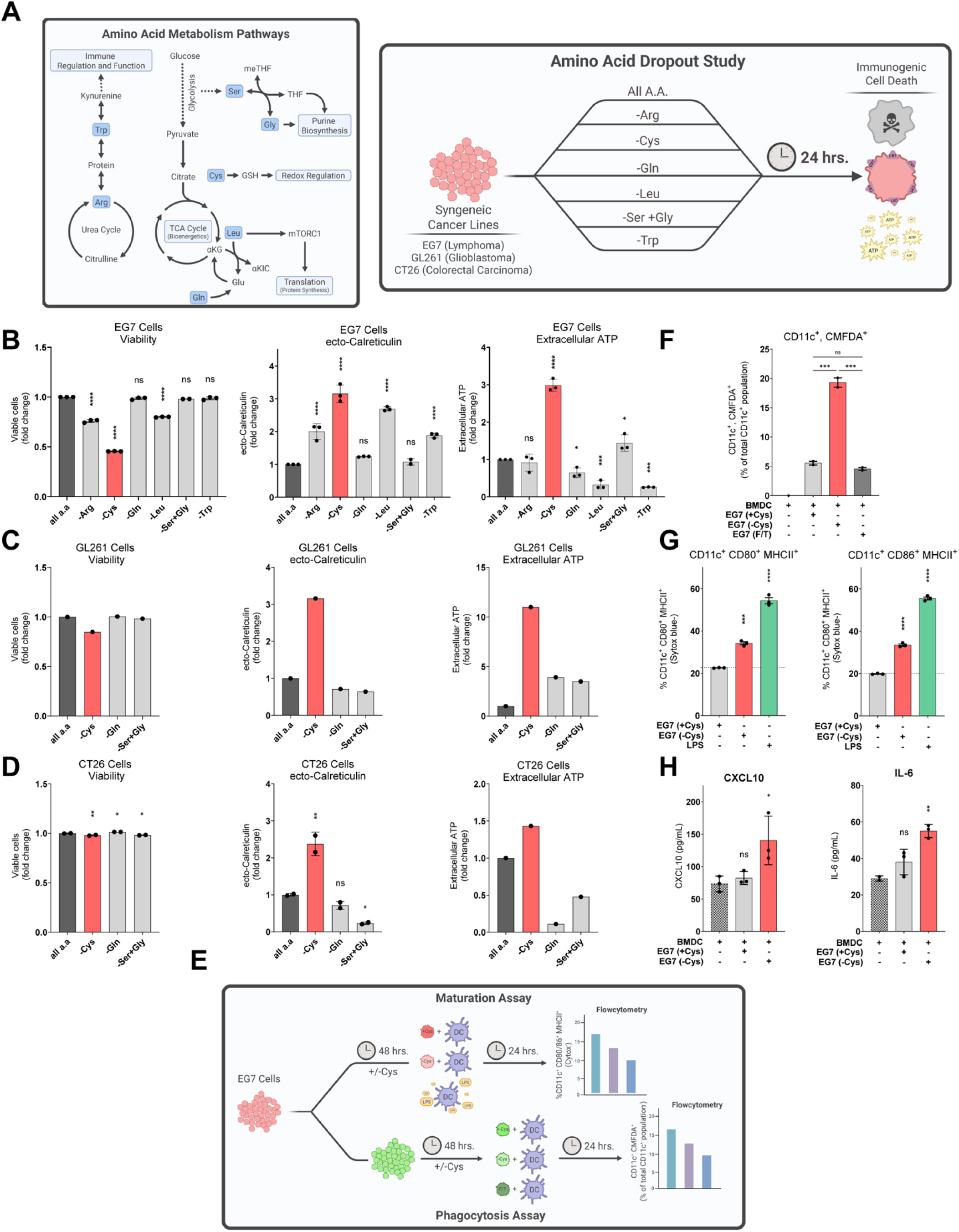
Cysteine depletion results in features of immunogenic cell death. **A**, Scheme of the amino acid metabolism pathways targeted in single amino acid dropout studies and the experimental setup of those studies. **B, C, D,** Viability, ecto-calreticulin, and extracellular ATP measured in EG7 (**B**), GL261 (**C**), and CT26 (**D**) cell lines at 24 hours under single amino acid dropout conditions. **E,** Experimental scheme depicting maturation and phagocytosis assays with murine BMDMs. **F,G,** Flow cytometry analysis of murine BMDCs assessing (**F**) phagocytosis of co-incubated EG7 cells, (**G**) as well as maturation markers CD80, CD86, and MHCII. EG7 pretreatment conditions: Full RPMI (+Cys), cystine-free RMPI (-Cys), and freeze/thaw (F/T). **H,** ELISA of supernatant from phagocytosis assay measuring CXCL10 and IL-6. Individual data points are presented over bar graphs with error bars, which represent the mean ± SD of technical replicates, where ns is not significant, *P* ≥ 0.05; *\*P* < 0.05; *\*\*P* < 0.01; *\*\*\*P* < 0.001; *\*\*\*\*P* < 0.0001.

From our assay media, we omitted tryptophan (Trp) and arginine (Arg), due to their involvement in immune regulation and function; glutamine (Gln) and Leucine (Leu), for their relationship to mTORC1 signaling and its downstream role in selective protein synthesis; serine (Ser) and glycine (Gly), for their roles in purine biosynthesis; and cysteine (Cys) for its importance in redox regulation. In these assays, EG7 cells were grown in media without each of the individual amino acids noted. After 24 hours, viability and DAMPs were measured. The latter included release of ATP and exposure of calreticulin. Cysteine restriction elicited the greatest release of ATP and calreticulin exposure among amino acid deprivation conditions (**Fig. 1B**). These results were further reflected in two additional syngeneic tumor models (CT26 and GL261), directing the focus of our studies on cysteine (**Fig. 1C,D**).

After measuring cell autonomous ICD features, we next sought to assess whether cysteine restriction in EG7 cells was sufficient to elicit immune stimulation. Dendritic cells sit at the crossroads of adaptive and innate immune responses and are required for antitumoral immune activity^68,69^. For effective antigen-cross presentation and T cell priming, both dendritic cell phagocytosis and maturation are essential^70–73^. Therefore, to assess these processes, we proceeded with co-culture studies of cysteine deprived EG7 cells and bone marrow derived dendritic cells (BMDC). In these studies, EG7 cells were grown in cysteine deprived or replete conditions for 48 hours prior to coculture with BMDCs (**Fig. 1E**). BMDC phagocytosis of EG7 cells was greater in co-cultures with cysteine deprived EG7 cells than in co-cultures with cysteine replete or freeze-thaw controls (**Fig. 1F, Supp Fig. 1A**). Free-thaw was used as an alternative, non-immunogenic form of cell death. Additionally, markers of BMDC maturation (CD80, CD86, and MHCII^74^) as well as production of proinflammatory cytokines (IL6 and CXCL10) were enhanced in co-cultures with cysteine deprived EG7 cells (**Fig. 1G,H**). Altogether, these data demonstrate the ability of cysteine deprived EG7 cells to stimulate BMDC functions of phagocytosis, maturation, and cytokine production.

### GPX4 inhibition-induced ferroptosis promotes features of immunogenic cell death in PDAC

Ferroptosis is a metabolic form of regulated cell death involving amino acid, iron, and lipid metabolism that results from uncontrollable lipid oxidation and cell rupture^75–83^. Prompted by our preliminary data with cysteine deprived EG7 cells, and the well appreciated role of cysteine in ferroptosis (**Fig. 2A**), we next compared cell death by ferroptosis inducers and cysteine restriction.

**Figure 2.**
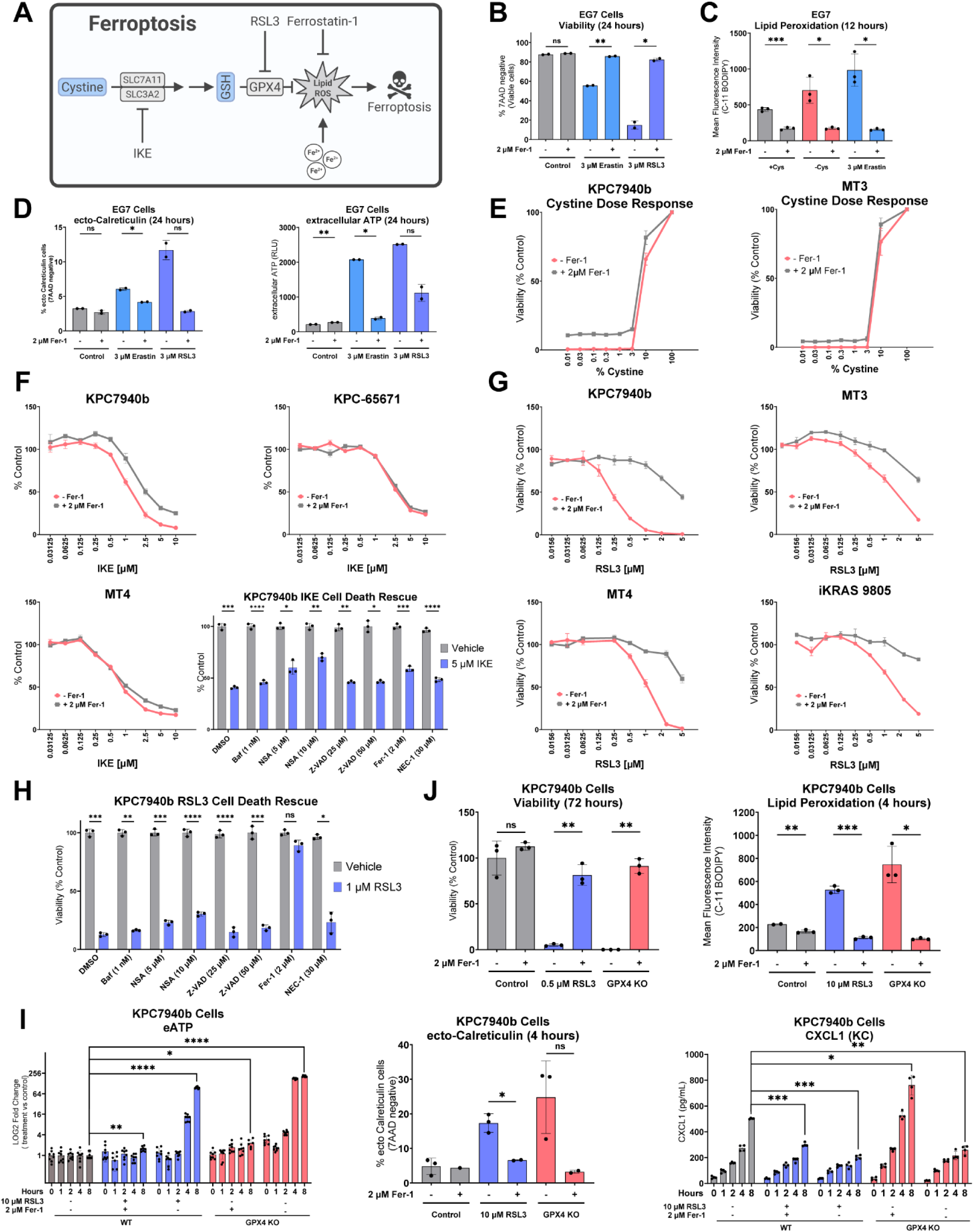
GPX inhibition-induced ferroptosis in PDAC cells results in features of immunogenic cell death. **A**, Scheme depicting canonical ferroptosis pathway. **B**, Viability of EG7 cells after 24-hour treatment with 3 µM RSL3 or 3 µM erastin, +/- 2 µM Fer-1. **C**, Lipid peroxidation of EG7 cells after 12-hour incubation with complete RPMI (+Cys), RPMI without Cystine (-Cys), or 3 µM erastin in complete RMPI, each condition +/- 2 µM Fer-1. **D**, ecto-Calreticulin positive EG7 cells and the ATP measured in their media after 24-hour treatment with 3 µM RSL3 or 3 µM erastin, each condition +/- 2 µM Fer-1. **E**, Viability of KPC7940b and MT3 cells grown under varied concentrations of cystine for 96 hours +/- 2 µM Fer-1, where [cystine] at 100% = 0.0652 g/L (comparable to complete RPMI). **F**, 48-hour dose response curves of imidazole ketone erastin (IKE) treated KPC7940b, KPC65671, MT4 cells +/- 2 µM Fer-1. (below) Viability of KPC7940b cells after 48-hours co-treatment with IKE and various cell death inhibitors. **G**, 48-hour dose response curves of RSL3 treated KPC7940b, MT3, MT4, and iKRAS 9805 cells +/- 2 µM Fer-1. **H**, Viability of KPC7940b cells after 48 hours of co-treatment with 1 µM RSL3 and various cell death inhibitors. **I**, (left) ATP and (right) CXCL1 measured in the media of 10 µM RSL3 treated or GPX4-deleted KPC7940b cells +/- 2 µM Fer-1 at time points from 0-8 hours. (middle) ecto-Calreticulin positive KPC7940b cells after 24-hour treatment with 10 µM RSL3 or withdrawal of 2 µM Fer-1 from GPX4-deleted KPC7940b cells. **J**, Viability of KPC7940b cells after 72-hour treatment with 0.5 µM RSL3 or after 72-hour withdrawal of 2 µM Fer-1 from GPX4-deleted KPC7940b cells. Lipid peroxidation measured in KPC7940b cells after 12-hour treatment with 10 µM RSL3 or 12-hour withdrawal of 2 µM Fer-1 from GPX4-deleted KPC7940b cells. Individual data points are presented over bar graphs with error bars, which represent the mean ± SD of technical replicates, where ns is not significant, *P* ≥ 0.05; *\*P* < 0.05; *\*\*P* < 0.01; *\*\*\*P* < 0.001; *\*\*\*\*P* < 0.0001.

Cystine restriction or inhibition of cysteine import are well established inducers of ferroptosis^84^. Indeed, EG7 cell death by cysteine depletion was rescued by the lipid peroxyradical trapping drug Ferrostatin-1 (Fer-1)^85^, a phenomenon similarly observed in cells treated with erastin, an inhibitor of cysteine import through system x ^- 86^ (**Fig. 2B, Supp Fig. 1B-C**). To further interrogate ferroptosis, we measured lipid reactive oxygen species (ROS), the ultimate mediator of ferroptosis. Comparable to erastin treated cells, cysteine deprived EG7 cells demonstrated lipid peroxidation that was rescued by Fer-1 (**Fig. 2C**). Consistent with our initial cysteine deprivation assays, 24-hour treatment with erastin induced DAMPs, extracellular ATP and ecto-calreticulin, in EG7 cells, and this was rescued by Fer-1 (**Fig. 2D, Supp Fig. 1D**). These results were also recreated by treatment with RSL3, an inhibitor of the antioxidant protein glutathione peroxidase 4 (GPX4) that prevents ferroptotic cell death by detoxifying oxidized phospholipids^77,83,87,88^ (**Fig. 2A-B, 2D**). Together, these results suggest that the immunogenic properties of cysteine deprivation result from ferroptotic cell death.

In syngeneic PDAC cells (KPC7940b, MT3, KPC-65671), cystine deprivation through media dropout or system x_C_^-^ inhibition with immidazone ketone erastin (IKE) leads to cell death; however, this was not meaningfully rescued by Fer-1 in vitro, suggesting a mechanism of cell death independent of ferroptosis (**Fig. 2E,F**). In contrast, RSL3-induced cell death was rescued by Fer-1 in four syngeneic murine PDAC cell lines (**Fig. 2G**). Furthermore, using a panel of autophagy, necrosis, apoptosis, or necroptosis inhibitors, only Fer-1 conferred a rescue of RSL3-induced cell death, reflecting ferroptosis as the mediator of RSL3-induced cell death in our PDAC model (**Fig. 2H**).

Thus, for the subsequent PDAC studies, we investigated the immunogenicity of GPX4-inhibition-induced ferroptosis. To this end, we used RSL3 alongside a GPX4 knockout cell line propagated in Fer-1 supplemented media (**Supp Fig. 1E**). We chose the KPC7940b cell model due to its sensitivity to RSL3-mediated GPX4 inhibition and robust rescue by Fer-1 (**Fig. 2G, Supp Fig. 1F,G**). Induction of ferroptosis by RSL3 treatment or restriction of Fer-1 from GPX4 knockout cells led to cell death and lipid peroxidation that could be rescued by Fer-1 (**Fig. 2J**). Importantly, induction of ferroptosis by RSL3 treatment or restriction of Fer-1 from GPX4 knockout cells also lead to the rapid release of ATP, exposure of ecto-calreticulin, and the secretion of CXCL1^61^ (**Fig. 2I**). The induction of these DAMPs was blocked when cells were co-treated with Fer-1.

Characteristics of ferroptotic cell death have been previously divided into three, time-dependent stages^61^. During initial ferroptosis (0-2 hours after the induction of ferroptosis), cells begin to accumulate lipid ROS. This is followed by an intermediate stage (3-4 hours), characterized by partial permeabilization of the cell membrane, leading to the release of ATP and the exposure of calreticulin. Finally, in terminal ferroptosis (6-8 hours), complete membrane permeabilization occurs, resulting in the release of LDH, HMGB1, and various cytokines. Our results demonstrating a temporal release of ATP, ecto-calreticulin exposure, and dampened CXCL1 secretion by ferroptotic cells coincided closely to these stages (**Fig. 2I**). Taken together, our data illustrate that ferroptosis initiated by GPX4 inhibition or knockout leads to features consistent with an immunogenic form of cell death.

### GPX4 inhibition-induced ferroptosis in PDAC cells leads to the selective release of immunosuppressive metabolites and lipids

Next, to identify metabolite-derived immunomodulatory signals, we set out to profile intra- and extracellular metabolomes and lipidomes by mass spectrometry. Informed by the time course described above, we selected a four-hour time point for the metabolomic and lipidomic analyses. Here, we employed two orthogonal models of ferroptosis: RSL3-treated wild type KPC7940b cells and Fer-1-withdrawn GPX4 knockout KPC7940b cells. We compared the relative abundance of individual metabolites in pellets and media supernatants of cells under basal and ferroptotic conditions. To ensure the identified metabolites were released from living cells undergoing ferroptosis, rather than cells whose plasma membrane has ruptured and their contents released, we analyzed extracellular levels of glycolytic intermediates across conditions as a surrogate for membrane rupture. Phosphorylated compounds are generally not secreted from cells, with exceptions for some transported metabolites (**Supp Fig. 1H-I**).

Media supernatants from RSL3-treated and GPX4 knockout cells revealed a consistent increase in the secretion of select nucleosides and nucleotides, whose abundance could be reversed when ferroptosis was inhibited with Fer-1 (**Fig 3A-D**). A broader range of similarly regulated metabolites were observed in cell pellets of RSL3-treated and GPX4 knockout cells, with a distinct signature suggesting nucleoside and nucleotide metabolism in ferroptotic cells (**Fig 3E-H**). Of the metabolites that we identified in extracellular fractions of ferroptotic cells, adenylates (e.g. ATP and AMP) are the most well reported for their ability to modulate immune activity^89–94^. Adenosine, which is well appreciated for its immunosuppressive activity^95–109^, can be derived from ATP and AMP in the TME through a series of conversions involving the ectonucleosides CD39 and CD73^110–116^. In addition, 2-Deoxyguanosine 5-monophosphate (dGMP) has been reported to suppress the secretion of IFN-γ from peripheral blood mononuclear cells when co-cultured with viral antigen^117^.

**Figure 3.**
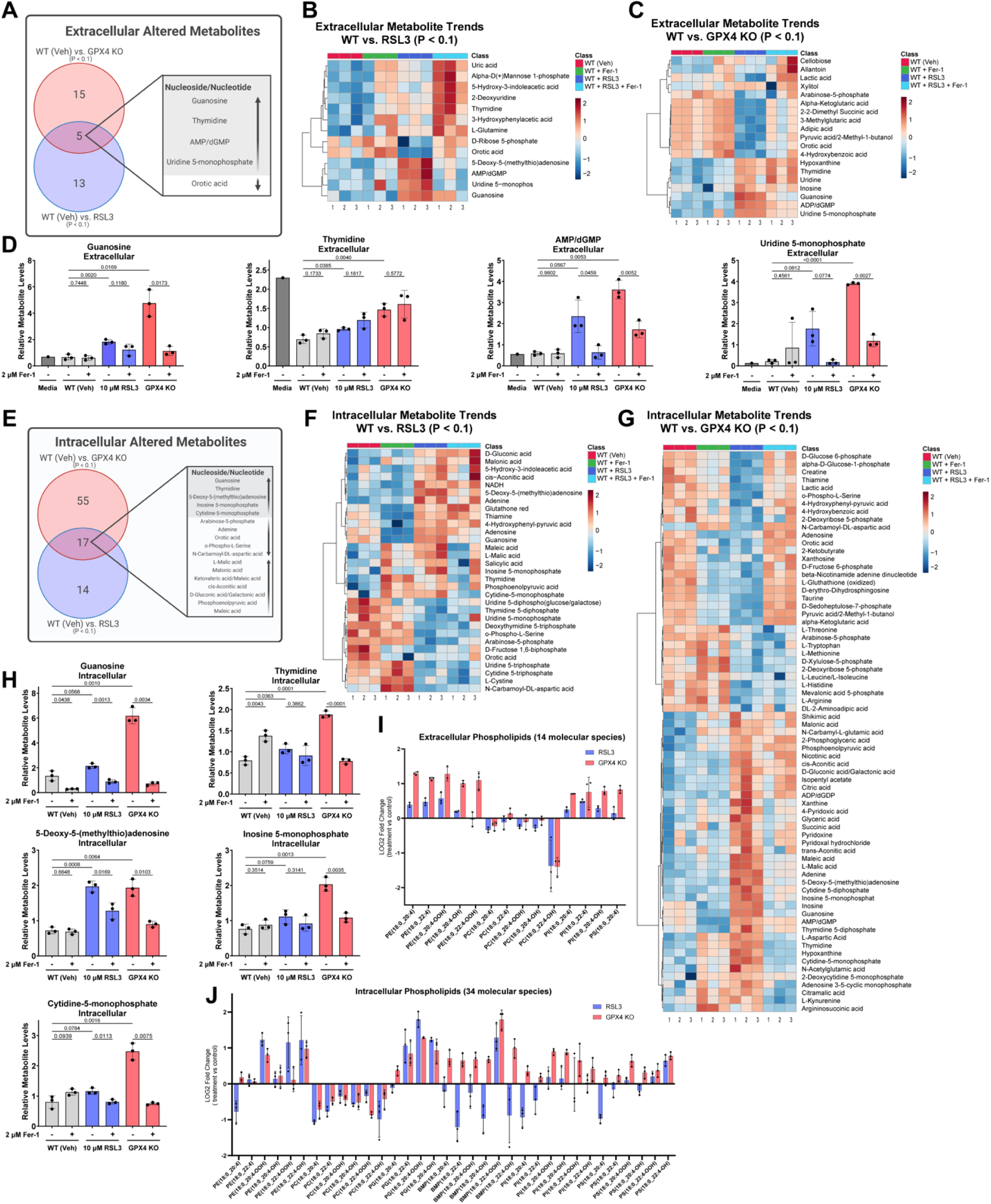
GPX inhibition-induced ferroptosis in PDAC cells results in the selective secretion of immunosuppressive metabolites. **A**, Scheme of metabolomics study showing consistently changing metabolites in the culture media of RSL3 treated and GPX4 KO KPC7940b cells, relative to vehicle. **B**, Heatmap showing differentially changed (P < 0.1) metabolites in the media supernatants of RSL3-treated and WT (Veh) KPC7940b cells **C**, or differentially changed (P < 0.1) metabolites in media supernatants of GPX4 KO and WT (Veh) KPC7940b cells. Log10 fold change is shown. **D**, Relative abundance of guanosine, thymine, Adenosine 5-monophosphate/2-Deoxyguanosine 5-monophosphate (AMP/dGMP), and Uridine 5-monophosphate in media supernatants of WT (Veh), RSL3-treated, and GPX4 KO KPC7940b cells, each with and without ferrostatin treatment. **E**, Scheme of metabolomics study showing consistently changing metabolites in the cell pellets of RSL3 treated or GPX4 KO KPC7940b cells. **F**, Heatmap showing differential (P < 0.1) metabolites in the cell pellets of RSL3-treated and WT (Veh) KPC7940b cells **G**, or differential (P < 0.1) metabolites in cell pellets of GPX4 KO and WT (Veh) KPC7940b cells. Log10 fold change is shown**. H**, Relative abundance of guanosine, thymidine, 5-Deoxy-5-(methylthio)adenosine, inosine 5-monophosphate, and cytidine-5-monophosphate in cell pellets of WT (Veh), RSL3-treated, and GPX4 KO KPC7940b cells, each with and without ferrostatin treatment. **I**, Log2 fold change of oxidized phospholipids detected by targeted mass spectrometry in supernatants of RSL3-treated and GPX4 KO KPC7940b cells as compared to WT (Veh) control. **J**, Oxidized phospholipids detected by targeted mass spectrometry in cell pellets of RSL3-treated and GPX4 KO KPC7940b cells as compared to WT (Veh) control. Individual data points are presented over bar graphs with error bars, which represent the mean ± SD of technical replicates, where ns is not significant, *P* ≥ 0.05; *\*P* < 0.05; *\*\*P* < 0.01; *\*\*\*P* < 0.001; *\*\*\*\*P* < 0.0001.

Next, we analyzed the intra and extracellular lipidomes of ferroptotic cells. Lipid peroxidation acts as a central driver of ferroptotic cell death, with the accumulation of oxidized polyunsaturated phospholipids leading to membrane damage, cell rupture, and ultimately cell death^118–120^. During ferroptosis, the most well-reported oxidized phospholipids are polyunsaturated phospholipids containing phosphatidylethanolamines (PEs), particularly those with arachidonic acid (AA, C20:4) and adrenic acid (AdA, C22:4) side chains^121–124^. In addition to oxidized PEs, certain oxidized phosphatidylcholines (PCs), such as 1-palmitoyl-2-(5-oxovaleroyl)-sn-glycero-3-phosphocholine (POVPC) and 1-palmitoyl-2-glutaryl-sn-glycero-3-phosphocholine (PGPC)^125^, have also been detected during ferroptosis and are known to contribute to membrane damage and cell death^126^. While oxidized forms of other phospholipids like phosphatidylserine (PS) and phosphatidylinositol (PI) have been observed, their roles are less well characterized compared to PE and PC species. Using a parallel scheme as to that of our metabolomic analysis, LC-MS lipidomic analysis focused on oxidized phospholipids revealed elevated levels of phospholipid hydroperoxide in both intra- and extracellular fractions of ferroptotic cells (**Fig. 3I,J, Supp Fig. 2A-B**). Untargeted LC-MS showed similar trends of oxidized phospholipids between RSL3-treated and GPX4 knockout cells in media supernatants (**Supp Fig. 2C-D, Supp Fig. 3A-D**). Taken in whole, these lipidomic data are consistent with previous reports and suggest that lipids are a potential source of immunomodulatory signals emitted by ferroptotic PDAC cells.

### Oxidized lipids inhibit CD4^+^ and CD8^+^ proliferation and effector function

We next assessed the effect of PDAC ferroptosis conditioned media on CD8+ T cell activation. Using conditioned media from KPC7940b GPX4 knockout cells undergoing intermediate ferroptosis, we observed that CD8+ T cell proliferation and late activation (as indicated by CD44 expression) were suppressed, phenotypes not observed in either wild type cells or GPX4 knockout cells cultured in Fer-1 (**Fig. 4A, Supp Fig. 4A**). Guided by our extracellular lipidomics data, we hypothesized that the inhibitory property of ferroptotic cell condition media was a consequence of oxidized lipids. To test this hypothesis, we assessed the impact of an oxidized phospholipids (OxPL) pool or arachidonic acid (AA) directly on CD4+ and CD8+ T cells. AA spontaneously oxidizes in culture, serving as a defined alternative to OxPL. Indeed, OxPL and AA both inhibited CD4+ and CD8+ T cell proliferation and CD8+ T cell effector molecule production (i.e. IFNγ and Granzyme B) (**Fig. 4B-D**). To directly test T cell functionality, we generated CD8+ T cells from OT-I mice, which recognize the ovalbumin antigen (OVA). By engineering MC38 syngeneic murine colon cancer cells to express OVA, we could test the ability of CD8+ T cells to kill cancer cells. Indeed, in this assay, pretreatment of T cells with OxPL or AA significantly diminished their ability to kill cancer cells (**Fig. 4E**). These data illustrate that, like proliferation and cytokine expression, CD8+ T cell functions are also impaired by oxidized lipids. Taken together, our evidence thus far revealed both immunogenic (i.e. DAMP release, antigen presenting cell activation) and immunosuppressive (i.e. oxidized metabolite and lipid release, suppression of T cells) features of ferroptotic cell death in PDAC cells, the sum of which can only be reliably interrogated in vivo.

**Figure 4.**
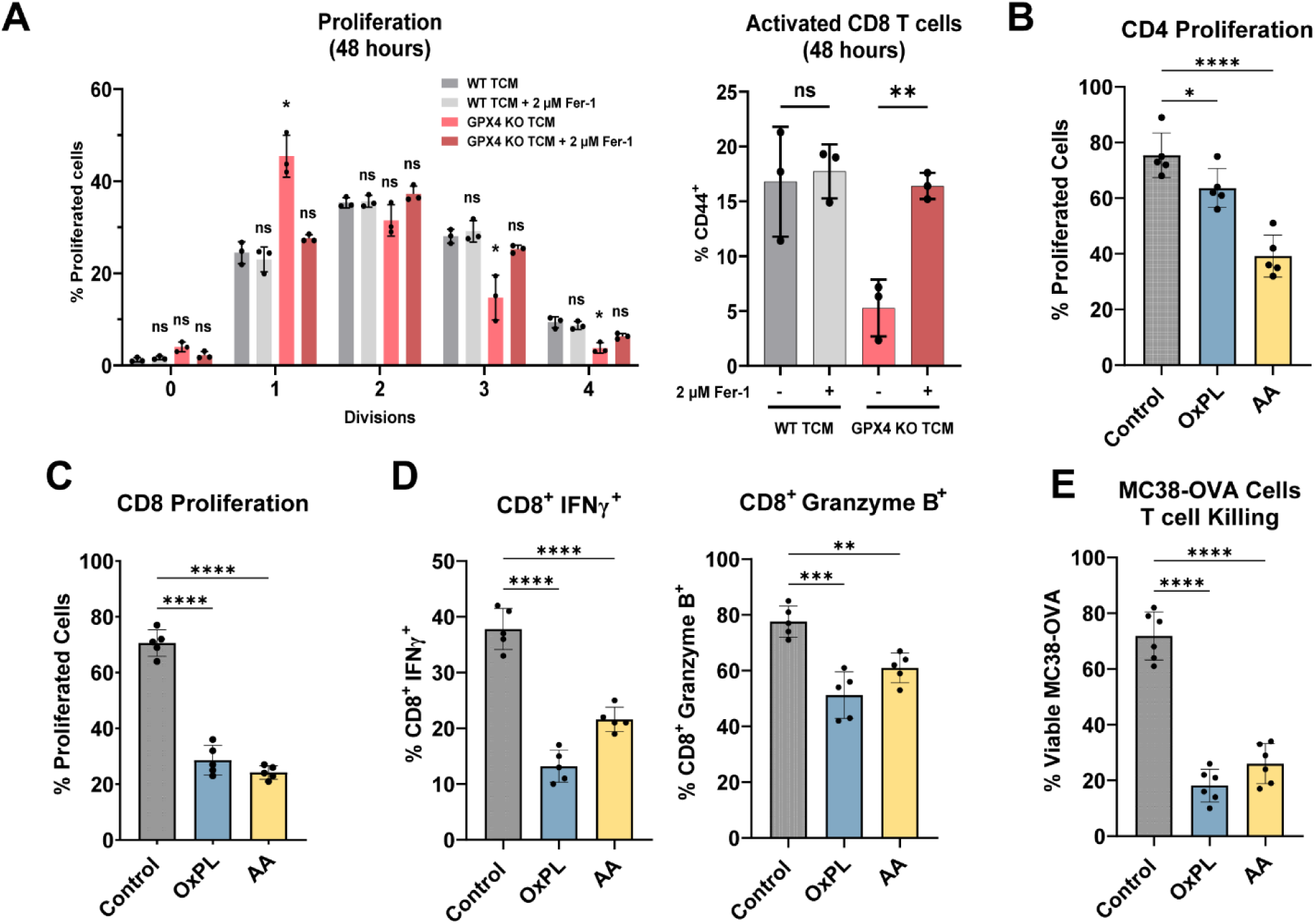
Oxidized lipids inhibit CD4^+^ and CD8^+^ proliferation and effector function. **A**, (left) Proliferation of CD8^+^ T cells cultured in conditioned media from WT KPC7940b cells grown in +/- 2 μM Fer-1 media or cultured in conditioned media from GPX4-deleted KPC7940b cells grown in +/- 2 μM Fer-1 media. (right) In parallel, the percentage of CD44^+^ CD8^+^ T cells were measured for these conditions. **B**, Proliferation of CD4^+^ and **C**, CD8^+^ T cells cultured in oxidized lipids (OxPL) or arachidonic acid (AA). **D**, Measurement of CD8^+^ IFNγ^+^ and CD8^+^ Granzyme B^+^ T cells cultured in OxPL or AA. **E,** T cell killing assay measuring the viability of MC38-OVA cells cocultured with OT-1 CD8^+^ T cells. OT-1 CD8^+^ T cells were preincubated in either OxPL or AA before coculture with MC38-OVA cells. Individual data points are presented over bar graphs with error bars, which represent the mean ± SD of technical replicates. Data in **D-E** represent technical replicates from two biologically independent samples. ns is not significant, *P* ≥ 0.05; *\*P* < 0.05; *\*\*P* < 0.01; *\*\*\*P* < 0.001; *\*\*\*\*P* < 0.0001.

### GPX4-inhibition promotes subcutaneous tumor outgrowth in tumor challenge experiments by altering immune populations

To determine how ferroptotic PDAC cell death influenced tumor immunity, we first attempted prophylactic vaccination using PDAC cells treated with an IC90 dose of RSL3 or positive control chemotherapies (i.e. mitoxanthrone or oxaliplatin), the gold standard for assessing ICD^41,42^. We observed tumor growth at the vaccination site uniquely for RSL3-treated cells (**Fig. 5A**); similarly prepared tumor cell vaccinations using chemotherapy treated cells did not result in tumor outgrowth at the vaccination site. These unexpected results suggested to us that ferroptosis, as opposed to chemotherapy-induced cell death, created a protective TME in vivo that enabled tumor outgrowth from the vaccine. In a subsequent study assessing whether our KPC7940b GPX4 knockout cells were a tractable option for prophylactic vaccination, we observed that PDAC cells lacking GPX4 could similarly establish and grow subcutaneous tumors (**Supp Fig. 4B-C**). This was in marked contrast to our results in vitro, where the presence of Fer-1 is required for these cells to remain viable (**Supp Fig. 1F**).

**Figure 5.**
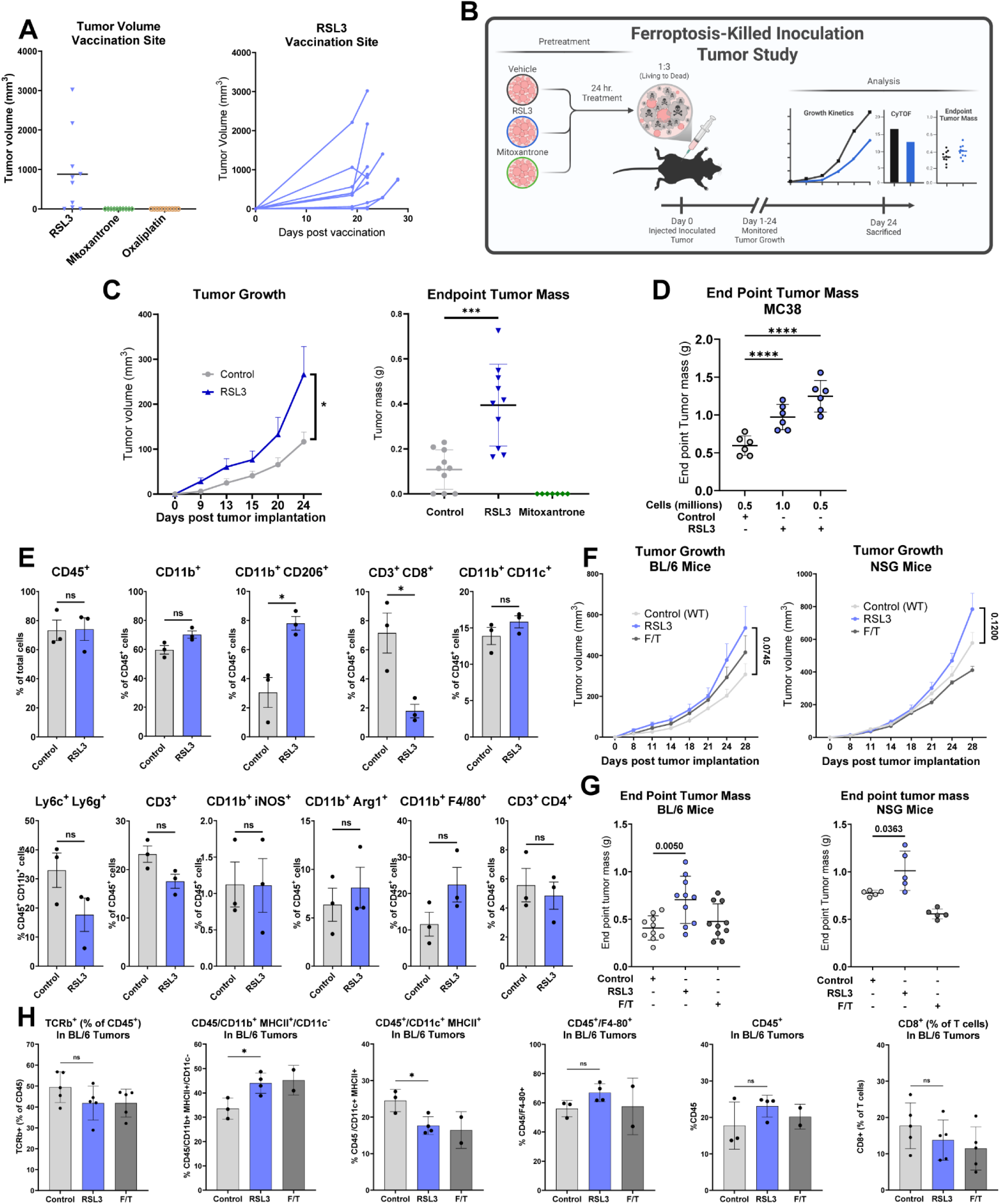
Inhibiting GPX4 in PDAC cells alters immune populations and enhances subcutaneous tumor outgrowth. **A**, (left) Endpoint subcutaneous tumor volume at vaccination site 22 days post prophylactic vaccination with 76% RSL3-killed, 71% mitoxantrone-killed, or 72% oxaliplatin-killed EG7 cells. Pre-injection cell viability was determined by flow cytometry. Mice vaccinated with mitoxantrone- and oxaliplatin-killed cells grew no tumors at their vaccination sites. All mice vaccinated with RSL3-killed cells were tumor-bearing at endpoint. (right) Subcutaneous tumor volume measured at the vaccination site in mice given RSL3-killed tumor vaccinations. **B**, Scheme of ferroptosis-killed tumor inoculation study with PDAC cells. **C**, (left) Subcutaneous tumor volume and (right) end point tumor mass of mice given a subcutaneous injection of living and RSL3-killed KPC7940b cells or a subcutaneous injection of an equivalent number of living cells. Tumors did not grow in mice given subcutaneous injections containing mitoxantrone-killed KPC7940b cells. **D**, End point tumor mass in BL/6 mice given subcutaneous injections of viable and RSL3-killed MC38 cells (RSL3) or viable MC38 cells alone (control). **E**, CyTOF analysis of subcutaneous tumors from three separate mice in RSL3 and Control arms from (**C**). **F**, Tumor volume measured across time **G**, and end point tumor mass in BL/6 and NSD mice given a subcutaneous injection of living and RSL3-killed KPC7940b cells (1:3) or living (RSL3) and freeze/thaw-killed KPC7940b cells (F/T) (1:3). Mice in the control arm (Control (WT)) were given a subcutaneous injection with an equivalent number of living KPC7940b cells. **H**, Flow cytometry analysis of subcutaneous tumors from separate mice in control, RSL3, and freeze/thaw tumor injection conditions from BL/6 mice in (**F,G**). Individual data points are presented over bar graphs with error bars, which represent the mean ± SD of technical replicates, where ns is not significant, *P* ≥ 0.05; *\*P* < 0.05; *\*\*P* < 0.01; *\*\*\*P* < 0.001; *\*\*\*\*P* < 0.0001.

To then study how ferroptotic PDAC cells influenced tumor growth, we inoculated wild type KPC7940b tumor injections with ferroptotic cells at a 1:3 living cell to dead cell ratio, and established subcutaneous tumors in immunocompetent C57BL/6 mice (**Fig. 5B**). Inoculation of subcutaneous tumors with mitoxantrone-killed wild type KPC7940b cells served as an immunogenic control, and indeed, this condition failed to produce tumors. Tumors inoculated with RSL3-killed KPC7940b cells were significantly larger in mass at endpoint and had increased growth kinetics compared to wild type KPC7940b tumors established with mock inoculations (**Fig. 5C**). Similarly, MC38 syngeneic colon cancer cells inoculated with RSL3-killed MC38 cells and then subcutaneously injected into syngeneic hosts grew significantly larger than MC38 cells injected alone (**Fig. 5D**). These data provide confirmatory results of the tumor growth promoting effects of ferroptotic cell debris in an independent cancer model.

Next, we used cytometry by time-of-flight (CyTOF) to evaluate the immune composition of the PDAC tumors in Figure 5C. The results of this analysis revealed that tumors inoculated with RSL3-killed cells were enriched with immunosuppressive myeloid cells (CD11b+/CD206+) and contained reduced populations of tumor infiltrating CD8+ T cells (**Fig. 5E, Supp Fig. 4D**). These data suggest that ferroptotic PDAC cells provided immune suppressive molecules that impaired the immune response, accelerating growth of PDAC tumors. To further assess the roles of the innate and adaptive immune compartments in GPX4-inhibited cancer cell immunosuppression, we implanted subcutaneous tumors inoculated with RSL3-killed KPC7940b cells in immune competent C57BL/6 mice (BL/6) or immunocompromised (NSG) mice (**Fig. 5F**). End point control tumors were significantly smaller than those inoculated with RSL3-killed cells in both genetic backgrounds (**Fig. 5G**). These results suggest that both the adaptive and the innate immune response potentiate tumor growth in the presence of ferroptotic cells. Similar trends of increased immune suppressive cells and decreased anti-tumor immune cells were observed in the ferroptotic arm of immune competent study, relative to control tumors (**Fig. 5H**).

## Discussion

In the present study, we document the production of adjuvants from cysteine deprived syngeneic murine cancer cell lines undergoing ferroptotic cell death, as well as their ability to induce maturation and enhanced phagocytosis of BMDCs. From this panel of murine models, we focused on PDAC. We demonstrate that induction of ferroptosis through RSL3 or genetic deletion of GPX4 results in *in vitro* features of ICD, which can be rescued by Fer-1. Our lipid-and metabolomic analysis of GPX4-inhibition-induced ferroptosis of PDAC cells showed the selective release of immunosuppressive metabolites and lipids, which may act as immune modulators in addition to canonical DAMPs. Indeed, we found that conditioned media impaired the proliferation, activation, and cytotoxicity of CD8+ T cells. To interrogate how these anti- and pro-tumor immune features integrate to influence tumor immunity, we analyzed subcutaneous PDAC tumors established with RSL3-killed PDAC or colon cancer cells. Results from these studies indicated an increase of immunosuppressive myeloid cells and diminished populations of tumor infiltrating CD8+ T cells, reflecting that pro-tumor ferroptotic signals outweigh the anti-tumor immune signals in our model systems. A subsequent study in immunocompetent BL/6 and immunodeficient NSG mice suggested that GPX4 inhibition in cancer cells suppresses both adaptive and innate immune responses. Taken together, these studies suggest that ferroptotic cell death aids in the establishment of a tumor protective milieu in syngeneic mouse models of PDAC.

While ferroptosis has emerged as an exciting new avenue for tumor therapy for PDAC^84,127–131^, careful investigation of its impacts on the tumor immune microenvironment, and overall consequence on tumor growth inhibition are necessary. Indeed, previous work from the field suggests that ferroptosis, under certain circumstances, may induce immune dysfunction and tumor progression. For example, phagocytosis of ferroptotic tumor cells by BMDCs has been shown to hamper their antigen-presenting capabilities and subsequently diminish T cell activation^61^. Ferroptotic tumor cells have also been found to release prostaglandin E₂^132^, which polarizes macrophages toward a tumor-supportive M2 phenotype and activates immunosuppressive cells like myeloid-derived suppressor cells and regulatory T cells^133,134^. In addition, lipid peroxidation products (e.g. 4-hydroxynonenal) generated during ferroptosis can also induce endoplasmic reticulum stress in immune cells, further exacerbating immune dysfunction^135^. Among a litany of others, these mechanisms illustrate how ferroptosis may facilitate tumor immune evasion and progression.

Conversely, T cell-driven ferroptosis has been shown to enhance the efficacy of immunotherapies like anti-PD-1 checkpoint blockade in preclinical models^136,137^. Clinical data also correlates markers of ferroptosis with positive responses to immunotherapy, suggesting that ferroptosis may play a role in the efficacy of immune checkpoint inhibitors^136^. In murine melanoma models, SLC7A11-deficient tumors displayed increased sensitivity to combined radiotherapy and anti-PD-L1 therapy, resulting in durable immunologic memory^138^. Another study showed that tumor cells sensitized to ferroptosis by inhibition of the itaconate transporter SLC13A3 enabled the effective treatment of immune checkpoint blockade-resistant tumors^139^. These seemingly inconsistent roles for the immunogenicity of ferroptosis highlight its complex role in modulating tumor-immune interactions.

While the presence of DAMPs and cytokines during cell death suggests ferroptosis could trigger ICD, the confluence of these features and their interaction with the TME introduces a myriad of additional considerations. For instance, ATP, a widely considered immunogenic DAMP, is readily converted to the immunosuppressive metabolite adenosine via CD39 and CD73 ectonucleosides that are expressed on tumor, stromal, and immune cells, making adenosine a prominent, immunosuppressive metabolite within the TME^140^. That is to say, while ferroptosis may endow dying tumor cells with adjuvant-like properties, our observations suggest that the TME imposes various mechanisms that neutralize the adjuvanticity of the DAMPs induced by ICD. Perhaps more importantly, our data supports the notion that DAMPs are not sufficient to predict the generation of anti-tumor immune responses. Our study instead illustrates the complexity signals released by dying tumor cells, such as DAMPs, metabolites, and lipids, and underlines the importance of in vivo studies to determine whether a mechanism of cell death is ultimately immunogenic or not.

In a recent study, Wiernicki et al. divided ferroptotic cell death into early, intermediate, and terminal stages, showing that DAMP release was primarily a feature of intermediate and terminal stages. Coincubation of early ferroptotic fibrosarcoma cells (i.e. syngeneic MCA205 cell line) hindered BMDC maturation, and phagocytosis, while, intermediate and terminal ferroptotic elicited robust maturation, and phagocytosis. Our study cooberates these findings, and extends them to PDAC. Further work suggested terminal ferroptotic MCA205 cells weakened the antigen presentation capabilities of BMDCs. As demonstrated herein and by others, high levels of oxPLs are produced during ferroptotis^141^ and may impair BMDC antigen presentation.^142–145^

Our metabolomics data showing nucleoside and nucleotide release is in agreement with recently published studies from Yapici, et al., who demonstrate similar findings in GPX4 knockout small cell lung carcinoma cells^132^. Their metabolomics profiling of cell pellets from inducible GPX4 knockout mouse embryoninc fibroblasts revealed a strong enrichment of purine and pyrimidine derivatives in cell pellets during early deletion of *Gpx4* before cell death was measured. In addition, they also observed the release of pyrimidine-related metabolites. Our data in syngenic murine PDAC cells do not fully align with their reported work in *Gpx4*-deleted MEFs; however, this may be attributed to differences in metabolic regulation, tissue of origin, or other factors disprate between murine PDAC cells and MEFs during ferorptosis-induced cell death. In another related study, Li, et al. showed that prophylactic vaccination with terminal ferroptotic cells confered protection against tumor challenges^131^. These cells were treated with the ferroptosis inducer N6F11 in vitro. In contrast, no significant protection was provided when mice were treated with a regiment of N6F11 in subcutaneous models, nor in orthotopic, syngineic allographs unless combined with anti-PD1 checkpoint therapy.

Like our work, other studies provide evidence suggesting that ferroptotic tumor cell death within the TME could result in adverse immune responses^146^. Li, et al., found that macrophage polarization states shift from anti- to protumor phenotypes under the influence of ferroptotic tissue damage, subsequently promoting *Kras*-driven PDAC tumorigenesis in mice^147^. Other groups have shown M2 macrophage polarization and myeloid-derived suppressor cell recruitment due to the liberation of KRAS and HMGB1 by ferroptotic cells in murine models of PDAC^147^ and mice with *Gpx4*-deficient liver tumors. Thus, the impact of these signals and other molecular mechanisms triggered by ferroptosis that are exclusively found in TME should be considered when evaluating the immunogenicity of ferroptosis, as their downstream effects may differ from those assessed peripherally through the prophylactic vaccination model. Further examination into both the tumor-intrinsic and extrinsic effects that dampen the adjuvanticity of ferroptotic cell death may uncover actionable targets and unlock the immunogenic potential of ferroptosis.

## List of abbreviations

PDAC: Pancreatic ductal adenocarcinoma
GPX4: glutathione peroxidase 4
Fer-1: Ferrostatin-1
TME: Tumor microenvironment
ATP: Adenosine triphosphate
DAMPs: Damage associated molecular patterns
ICD: Immunogenic cell death
BMDC: Bone marrow derived dendritic cell

## Declarations

### Ethics Approval

Animal experiments were approved by the University of Michigan Institutional Animal Care and Use Committee following PRO00010606.

### Consent for publication

Not applicable

### Availability of data and materials

All data generated or analyzed during this study are included in this published article and its supplementary information files.

### Author’s contributions

N.E.M., H.S.H., C.A.L., designed the study. N.E.M., H.S.H., D.S., C.A.L. designed experiments and N.E.M., H.S.H., R.E.M., R.S. performed the bulk of the experimental studies with support from M.P., J.A., A.C.A, L.Z., D.S. N.E.M., H.S.H., R.E.M., P.S., D.S. contributed to data analysis. C.A.L. provided resources, funding and conceptual input for experiments and supervised the research. D.S. and C.A.L. wrote the manuscript. D.S. curated data and prepared the manuscript figures. Other listed authors contributed crucial feedback and support. All authors reviewed and approved the final manuscript.

## Acknowledgments

We would like to thank Joel Whitfield and the Rogel Cancer Center Immune Monitoring Shared Resource Core team for assistance with cytokine assays, as well as Jeff Johnson, Paul Kennedy, and Randy Strube from Cayman Chemical for conducting LC-MS Lipidomics.

---

**Supplemental Figure 1.**
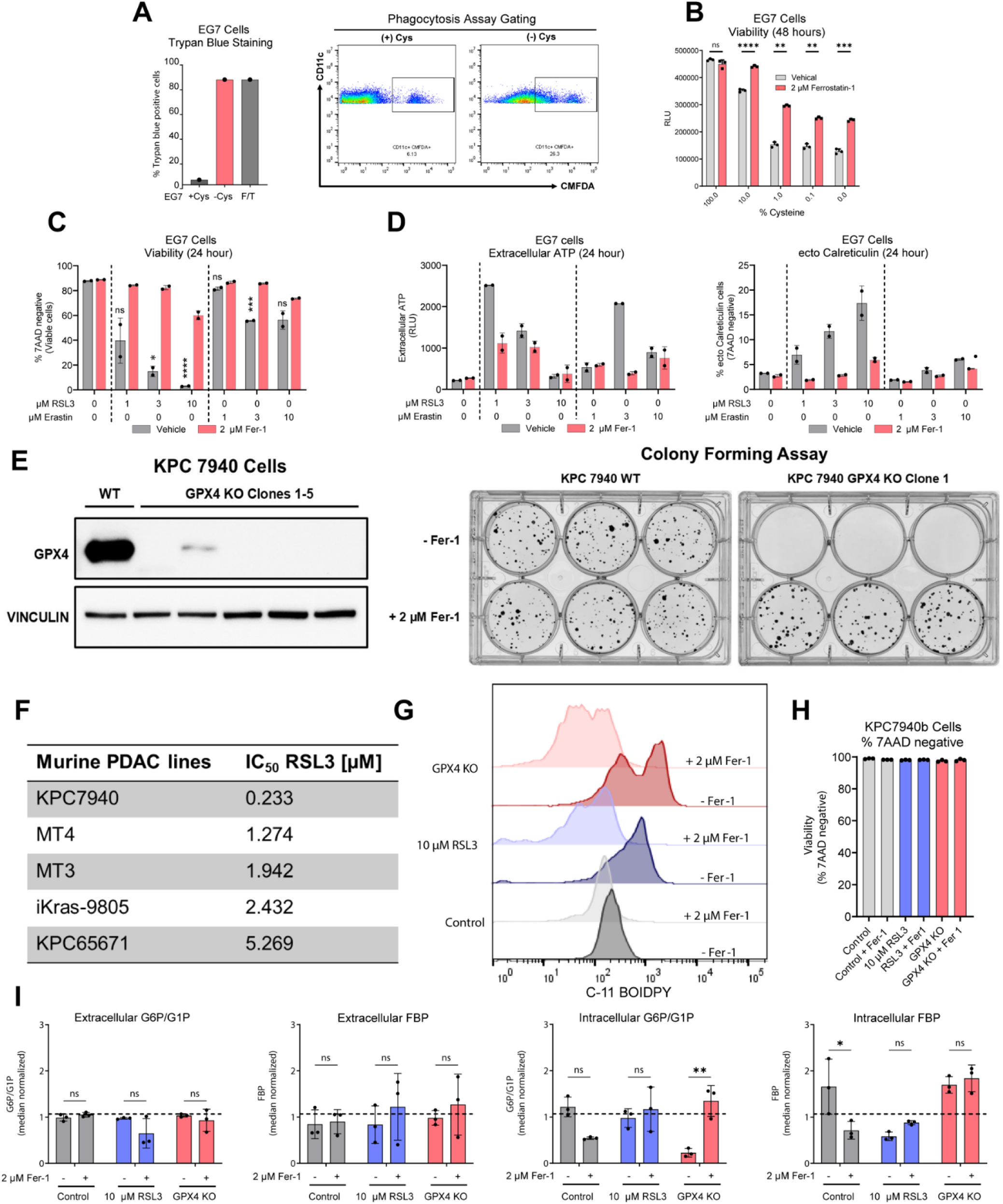
Analyses of cystine-deprivation and RSL3-treatment in murine cancer cell lines. **A**, (left) Trypan Blue staining of the EG7 cells used in phagocytosis assay. EG7 cells were incubated for 48 hours in complete RPMI (+Cys), RPMI without cystine (-Cys), or subjected to freezing-thawing (F/T). (right) Flow cytometry gating of CD11^+^ CMFDA^+^ cells in phagocytosis assay. **B**, Cell viability measured using CellTiter-Glo Luminescent Cell Viability Assay, presented in relative light units (RLU), of EG7 cells grown under varied concentrations of cystine for 48 hours with or without 2 µM Fer-1, where [cystine] at 100% = 0.0652 g/L (comparable to complete RPMI). **c**, Cell viability measured by 7-AAD of EG7 cells after 24-hour treatment with 1, 3, or 10 μM RSL3 or Erastin, each dose with and without 2 μM Fer-1. **D**, (left) Extracellular ATP measured by luminescence in supernatants of EG7 cells and (right) ecto-calreticulin of 7-AAD negative EG7 cells after 24-hour treatment with 1, 3, or 10 μM RSL3 or Erastin, each dose with or without 2 μM Fer-1. **E**, (left) Western blot of KPC7940b GPK4 KO single cell clones. (right) Six-day colony forming assay of KPC7940b WT and KPC7940b GPX4 KO clone 1 cultured with and without 2 μM Fer-1. **F**, RSL3 IC_50_ concentrations in a panel of murine PDAC cell lines. **G**, Lipid peroxidation measured by C-11 BODIPY in KPC7940b after 12-hour treatment with 10 µM RSL3 or 12-hour withdrawal of 2 µM Fer-1 from GPX4-deleted KPC 7940 cells. **H**, Viability of KPC7940b cells used in metabolomics as measured by 7-AAD. **I**, Mass spectrometry analysis of glucose-phosphate or fructose 1,6-bisphosphate detected in KPC 7940 cells treated with or without 10 μM RSL3 or GPX4 KO cells cultured with or without 2 μM Fer-1 for 4 hours. Individual data points are presented over bar graphs with error bars, which represent the mean ± SD of technical replicates, where ns is not significant, *P* ≥ 0.05; *\*P* < 0.05; *\*\*P* < 0.01; *\*\*\*P* < 0.001; *\*\*\*\*P* < 0.0001.

**Supplemental Figure 2.**
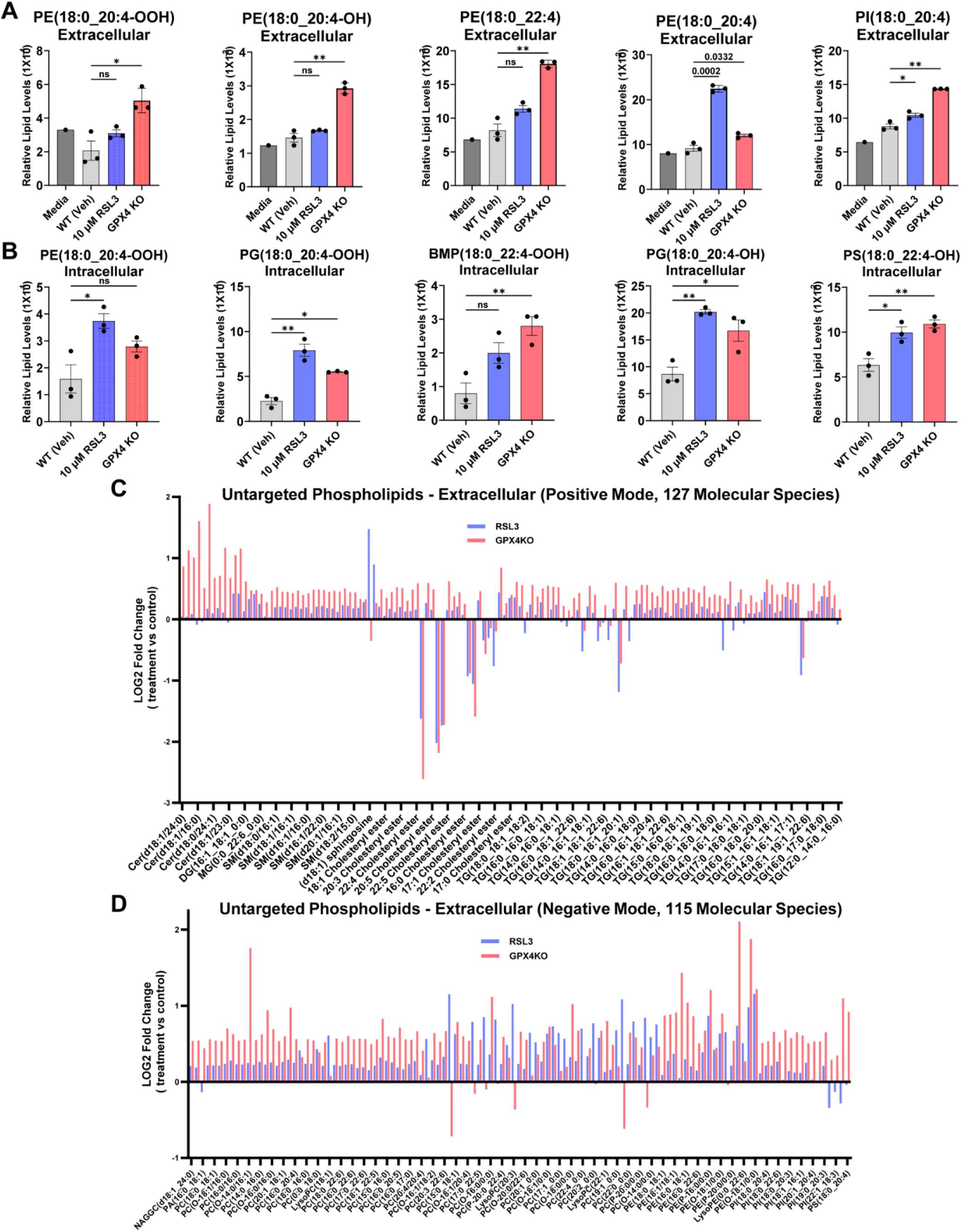
Targeted lipidomics analysis of ferroptotic PDAC cells. **A**, Panel showing relative lipid levels in cell-free media, and supernatants of WT (Veh), RSL3-treated, and GPX4 KO KPC 7940 cells detected by targeted mass spectrometry. **B**, Panel showing relative lipid levels in cell pellets of WT (Veh), RSL3-treated, and GPX4 KO KPC7940b cells. **C**, Log2 fold change of oxidized phospholipids detected by untargeted mass spectrometry (positive ion mode) in extracellular samples from RSL3-treated and GPX4 KO KPC7940b cells as compared to WT (Veh) control. **D**, Log2 fold change of oxidized phospholipids detected by untargeted mass spectrometry (negative ion mode) in extracellular samples of RSL3-treated and GPX4 KO KPC 7940 cells as compared to WT (Veh) control. All data are presented as the mean ± SD of technical replicates, where ns, not significant, *P* ≥ 0.05; *\*P* < 0.05; *\*\*P* < 0.01; *\*\*\*P* < 0.001; *\*\*\*\*P* < 0.0001.

**Supplemental Figure 3.**
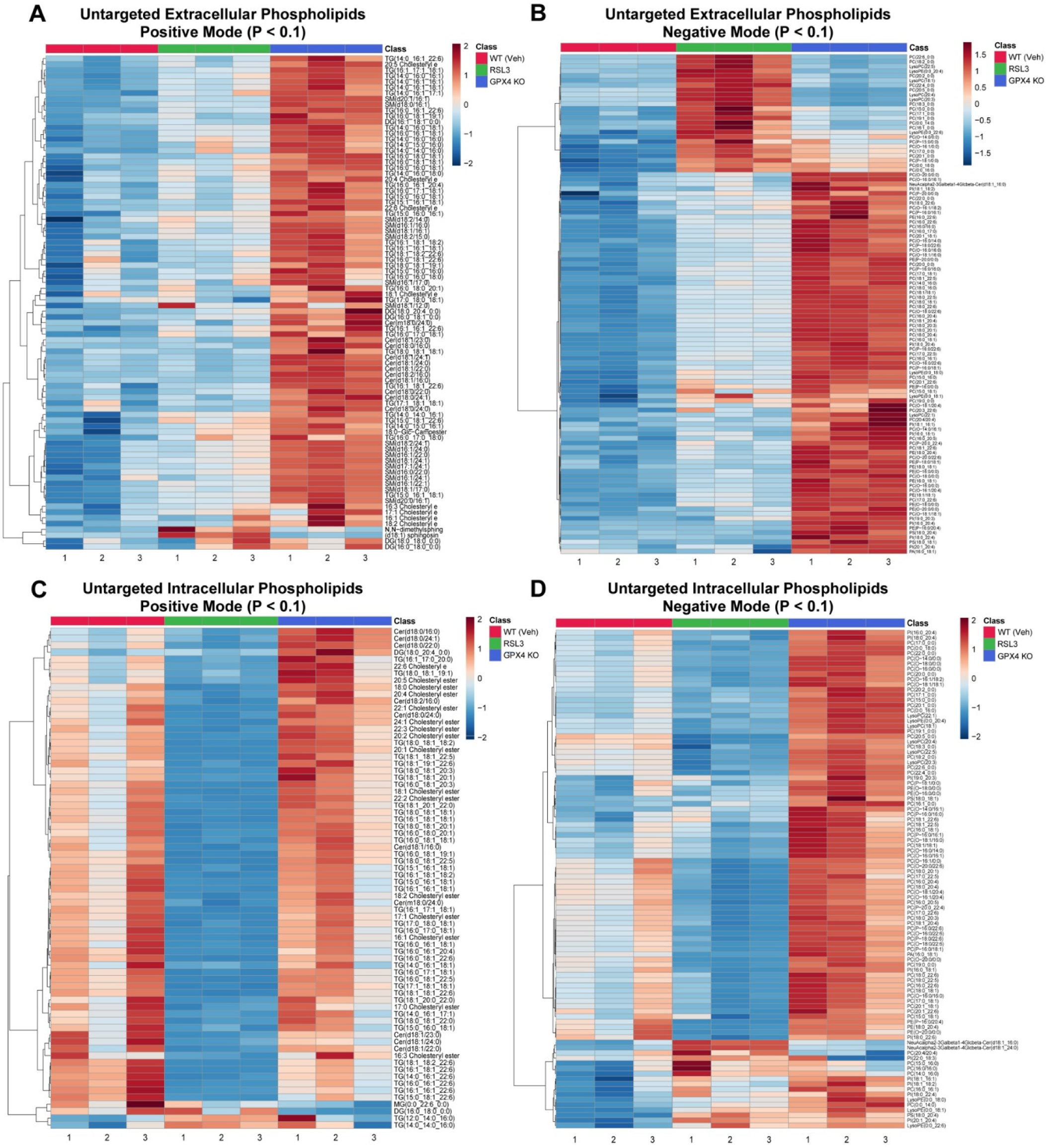
Heatmaps of lipidomics analysis in ferroptotic PDAC cells. **A,B**, Heatmap showing differential (P < 0.1) oxidized phospholipids detected by untargeted mass spectrometry in positive ion mode (**A**) or in negative ion mode (**B**) in extracellular extracts of RSL3-treated and GPX4 KO KPC7940b cells as compared to WT (Veh) control. Log10 fold change is shown. **C,D,** Heatmap showing differentiatl (P < 0.1) oxidized phospholipids detected by untargeted mass spectrometry in positive ion mode (**C**) or in negative ion mode (**D**) in cell pellets of RSL3-treated and GPX4 KO KPC7940b cells as compared to WT (Veh) control. Log10 fold change is shown.

**Supplemental Figure 4.**
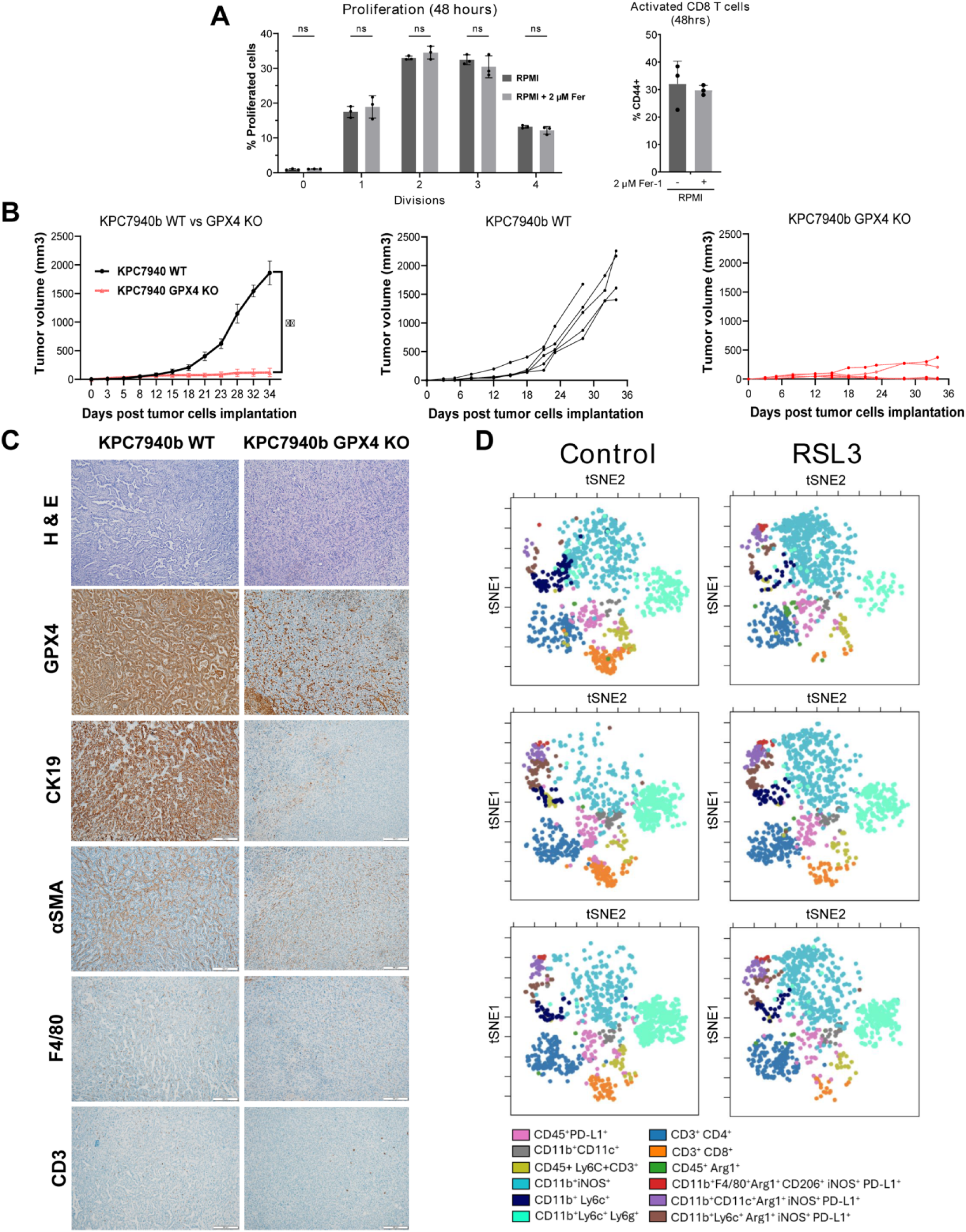
T cell and tumor analyses for the effects of PDAC ferroptotis. **A**, (left) Proliferation of CD8+ T cells cultured in complete RPMI with or without 2 μM Fer-1, measured by flow cytometry using Cell Tracker. (right) CD44+ CD8+ T cells were measured for these conditions. Data are presented as the mean ± SD of technical replicates. **B**, Subcutaneous tumor volume in KPC7940b WT and KPC7940b GPX4 KO tumors, plotted as groups (left) and by individual mice (middle, right). Data are presented as the mean ± SD of biologically independent samples. **C**, Representative immunohistology staining of KPC7940b WT and GPX4 KO tumors from (**B**). **D**, CyTOF analysis of three subcutaneous tumors from three separate mice in the RSL3-killed group and control group from Fig. 5C. Mice in the mitoxantrone condition did not develop tumors; therefore, CyTOF analysis could not be completed for this treatment condition. ns, not significant, *P* ≥ 0.05; *\*P* < 0.05; *\*\*P* < 0.01; *\*\*\*P* < 0.001; *\*\*\*\*P* < 0.0001.

**Supplementary File 1**. Raw metabolomics data used for the presentation of data in Fig. 3 and Supplementary Fig. 1I.

**Supplementary File 2**. Raw targeted lipidomics data used for the presentation of data in Fig. 3 and Supplementary Fig. 2.

**Supplementary File 3**. Raw untargeted lipidomics data used for the presentation of data in Fig. 3 and Supplementary Fig. 2.

**Supplementary File 4**. Unprocessed western blot images corresponding to the data from Supplementary Figure 1E.

